# neuroWalknet, a controller for hexapod walking allowing for context dependent behavior

**DOI:** 10.1101/2022.04.27.489633

**Authors:** Malte Schilling, Holk Cruse

**Author notes:** CORRESPONDING AUTHOR MAIL.

## Abstract

Decentralized control has been established as a key control principle in insect walking and has been successfully leveraged to account for a wide range of walking behaviors in the proposed neuroWalknet architecture. This controller allows for walking patterns with different velocities in forward and backward direction — quite similar to the behavior shown in stick insects —, for negotiation of curves, and for robustly dealing with various disturbances.

While these simulations focus on the cooperation of different, decentrally controlled legs, here we consider a set of biological experiments not yet been tested by neuroWalknet, that focus on the function of the individual leg and are context dependent. These intraleg studies deal with four groups of interjoint reflexes. The reflexes are elicited by either a stimulation of the femoral chordotonal organ (fCO) or a specific stimulation of campaniform sensilla (CS). Motor output signals are recorded from the alpha-joint, the beta-joint or the gamma-joint of the leg. Furthermore, such reflexes have been studied while the ganglion was treated with pilocarpine.

Although these biological data represent results obtained from different local reflexes in different contexts, they fit with and are embedded into the behavior shown by the global structure of neuroWalknet. In particular, a specific and intensively studied behavior, active reaction, has since long been assumed to represent a separate behavioral element, from which it is not clear why it occurs in some situations, but not in others. This question could now be explained as an emergent property of the holistic structure of neuroWalknet. When experimenting with pilocarpine, oscillations were induced in neuroWalknet even though this does not include an explicit central pattern generator and in this way provides a simpler model as a functional explanation. As the simulation data result from a holistic system, further results were obtained that could be used as predictions to be tested in further biological experiments.

**AUTHOR SUMMARY:** Behavior of animals can be studied by detailed observation, but observation alone does not explain the function of the underlying neuronal controller structures. To better understand this function, an important tool can be to develop an artificial structure based on simulated neurons and a simulated or physical body. Although typical animal behavior appears complex, the corresponding neuronal structures may be comparatively simple.

The goal for such a hypothetical structure should be to include as many different behaviors as possible, and, at the same time, search for a simple explanation consisting of a minimum of neuronal elements. Furthermore, such a simulation system, e.g. an artificial neuronal network, should contain hypotheses that can be tested in biological experiments.

We propose an extension to such a network that is based on a decentralized neuronal structure, using a neural network as a scaffold, that enables various combinations of local neuronal elements that allow for emergent, i.e. not explicitly designed properties. Indeed, neuroWalknet contains further abilities not yet recognized in the earlier version. For instance, neither explicit structures like central pattern generators nor explicit Active Reaction are required to reproduce typical intraleg reactions. Therefore, neuroWalknet presents a holistic approach enabling emergent properties out of the cooperation of small neuronal elements that are context dependent instead of explicit, dedicated elements.

## 1 Introduction

Animals are able to perform a large variety of complex behaviors, which makes it a challenging task to unravel the underlying control structures. As a first difficulty, when visually observing different behaviors, it is usually not clear if these behaviors or parts of the behaviors are controlled by separate neuronal structures or result from a combination of various small elements being activated due to the current context. Furthermore, as a second problem, structures functionally defined in one context may not show the same function in another context. Therefore, we have to address these questions as to (i) how could possible functional elements be defined and (ii) how may they be combined to allow for control of complex adaptive behavior?

Simulation studies in combination with biological behavior and neuronal studies offer a sensible tool to answer the second question as to how hypothetical behavioral elements might be combined to form minimal (e.g. mechanical or neuronal) structures that may explain specific behaviors. Concerning the first question, we will start using a functional control structure proposed earlier (Schilling & Cruse, 2020). The focus of this controller has been on hexapod walking which represents a basic but already complex behavior as it deals with many degrees of freedom. Therefore, such a system constitutes an example well suited to study the principles of how animal behavior might be controlled. In this article, it is our goal to further analyze how the proposed structures can explain additional behavioral findings which would then offer further credence to this hypothetical structure and the underlying principles of that controller.

This network, named neuroWalknet (Schilling & Cruse, 2020), together with robot Hector (Dürr et al., 2019), followed this line of testing hypothetical mechanisms in simulations. Indeed, it allows to control the basic different aspects of walking as are forward walking and backward walking, negotiation of curves, different velocities, walking on irregular ground, climbing, but also those concerning activation of partial or complete deafferentation of legs. In other words, neuroWalknet focuses on interleg control and was able to demonstrate that the functional elements of this controller can account for all these behaviors. The different behaviors emerge out of the interaction between the modules of the control system and with the environment. As such, neuroWalknet has proven as a strong candidate system that can already deal with a large variety of complex behavior and that captures important underlying control principles.

There are however a number of experimental results focusing on control of the individual leg — eventually addressed as “interjoint reflexes” —, for which no common detailed structure has been developed yet. This stimulates the question to what extent the already existing controller, neuroWalknet, may predict these results, too. If neuroWalknet can explain these experimental results as well, this would further support it as a trustworthy model. To tackle this challenge, in this article we will focus on experiments on intraleg control and will therefore address activation patterns of the three most relevant joints of the legs, in neuroWalknet called alpha joint (i.e. the Thorax-Coxa joint (T-C joint), the beta joint (i.e. Coxa-Trochanterofemur joint (C-T joint) and the gamma joint (i.e. Femur-Tibia joint (F-T joint). The corresponding muscle pairs are called Protractor and Retractor, Levator and Depressor, and Flexor and Extensor, respectively. Specifically, the focus will be on context dependency of reactions. In all addressed cases—all referring to stick insect studies—context dependent reflex changes will be treated that depend on the state of the animal, for example forward vs. backward walking or state swing vs. state stance.

Overall, we are considering four groups of experiments (Table 1). The first three experiments deal with reflexes on the single joint level and will in particular focus on context dependency. The first group of experiments studies reflexes in the alpha joint. Here, as an example for the type of experiments addressed, we will consider stimulation of campaniform sensilla (CS) by bending the femur to stimulate specific CS. In forward walking, caudal bending stimulates the retractor muscles in the alpha joint (Akay et al., 2004, 2007; Schmitz, 1993; Schmitz & Stein, 2000). In contrast, during backward walking as a different context and state, the same stimulus now activates the protractor. A context dependent reflex is observed, in this case a reflex reversal depending on the walking direction. The second and third group of experiments will deal with reflexes triggered by activation of the femoral Chordotonal organ (fCO) relevant for position and velocity of the gamma joint and the response in the gamma joint and beta joint, respectively.

**Table 1:** Overview of performed experiments and summary of the biological experiments.

| Exp. | Stimulus | Biol. Output | Joint | Exp. Var. | State | Observation | Explanation | Res. Fig. | Biol. Ref. |
| --- | --- | --- | --- | --- | --- | --- | --- | --- | --- |
| 1 | Campaniform Sensilla | Protractor-Retractor | Alpha | Direction | forward | caudal bending → retractor<br>rostral bending → protractor |  | Fig. 2 | Schmitz 1993, Schmitz & Stein 2000, Akay et al. 2004, 2007, Haberkorn et al. 2019) |
|  |  |  |  |  | backward | caudal bending → protractor<br>rostral bending → retractor | reflex reversal dep. on walking dir. |  |  |
| 2.1 | femoral Chordotonal Organ | Flexor-Extensor | Gamma | Curve Walking | inner leg | flexion of fCO → flexor | Active Reaction | Fig. 3 | Hellekes et al., 2012 |
|  |  |  |  |  | outer leg | flexion of fCO → const. or ext. | No Active Reaction |  |  |
| 2.2 | femoral Chordotonal Organ | Flexor-Extensor | Gamma | Direction | forward | flexion of fCO → flexor | Active Reaction | Fig. 4 | Hellekes et al., 2012, Bässler, 1986 |
|  |  |  |  |  | backward | flexion of fCO → extensor | reversal |  |  |
| 3 | femoral Chordotonal Organ | Levator-Depressor | Beta | Direction | forward | flexion of fCO → levator | No reflex reversal | Fig. 5 | Hess & Büschges, 1999, Bucher et al., 2003 |
|  |  |  |  |  | backward | flexion of fCO → levator |  |  |  |
| 4.1 | Campaniform Sensilla | Protractor-Retractor | Alpha | application of pilocarpine plus add single sensory stimulus (other connections cut) |  | caudal bending → entrain motor output |  | Fig. 7 | Akay et al., 2007 |
| 4.2 | femoral Chordotonal Organ | Flexor-Extensor | Beta |  |  | flexion of fCO → levator |  | Fig. 8 | Hess & Büschges, 1999 |

In all these experiments, the response depends on the context, in some cases producing opposite reflexes or showing a reversal of a reflex. Therefore, it appears important to consider the current state and context of the control system when interpreting reactions to stimuli. Further, we want to consider a fourth group of experiments in which the behavior of a (more or less) intact animal has been compared with animals experiencing the same treatment but, in addition, being treated with pilocarpine, a procedure that elicits oscillations in all three joints of a leg (Büschges et al., 1995). A common assumption is that quasi rhythmic leg movements required for walking are basically controlled by local joint oscillators (“central pattern generators”, CPG) plus sensory feedback for fine tuning, a view which has been considered as to be supported by the biological experiments addressed in the fourth group. As these experiments focus on single joint mechanism, we want to address the question whether the—in these experiments artificially induced—neuronal oscillations represent dedicated and necessary control structures that are active in the intact legs, too. In contrast, we show, as in the earlier study (Schilling & Cruse, 2020), that another explanation is possible which means that the experiments addressed here may not support the idea that CPG structures are necessary for control of walking.

In this article, we will test to what extent these (and related) experimental results can be described by neuroWalknet as an artificial neural network that has already been shown to simulate a broad range of behaviors (Schilling & Cruse, 2020). In this way, we will test if neuroWalknet could serve as a predictor for the biological studies. We will show that, with application of only a minor (technical) expansion, neuroWalknet will indeed explain the experimental results as discussed here. In addition, these simulation results contain further predictions that may be tested in new biological experiments. While in the biological experiments usually only motor output of one leg has been recorded, the simulation always produces outputs for all three leg joints which closes the loop from simulation study back to neurobiology and provides testable hypotheses for future experiments. Furthermore, concerning the contribution of CPGs to walking, as postulated by earlier authors, we show that a dedicated CPG structure might not be necessarily required which is in agreement with the earlier study (Schilling & Cruse, 2020). Following the notion that a less complex solution should be preferred, we propose to prefer this simpler solution that, in addition, can explain more biological data.

Thus, these results support—due to its predictive properties—the idea that neuroWalknet (Schilling & Cruse, 2020) represents a powerful hypothesis which is based on a simple functional structure that may represent a basic motor control system for insects in general.

## 2 Overview of extension of neuroWalknet – A structure for establishing different context

neuroWalknet has been realized as an artificial neural network that acts as a controller for a six-legged robot (Schilling & Cruse, 2020). Initially, it has been envisioned for control of locomotion for the case of forward and backward walking, but since then has been tested in a variety of behavioral contexts. The main governing principle is the idea of decentralization. While most control approaches employ one holistic control unit that controls all available degrees of freedom, in this decentralized approach control is distributed into different control units that are only loosely coupled. On the one hand, this helps simplify the control as the number of explicitly coordinated degrees of freedom is kept low per controller. On the other hand, still it has shown that stable behavior emerges in different contexts and the approach became more robust to perturbations from the environment. Decentralization of control is realized through individual controllers for each leg. This type of decentralization is shared with the family of earlier Walknet approaches (Cruse et al., 1998; Dürr et al., 2004; Schilling, Paskarbeit, et al., 2013; Schilling, Hoinville, et al., 2013). Each leg is controlled by a single controller which decides on which action should be performed by a leg. Different actions are represented by so called Motivation Units, simple artificial neurons with partially linear characteristic and low-pass filter dynamics (Fig. 1, grey units). There is a competition between these units on the leg level which is sensory driven (e.g., a swing movement is ended when ground contact is established). Such decentralized structures have been widely used for simulation of adaptive walking behavior and have been extended towards planning control architectures (Schilling, Paskarbeit, et al., 2021; Schilling & Cruse, 2017) as well as learning-based approaches (Schilling et al., 2020; Schilling, Melnik, et al., 2021; Schilling & Melnik, 2018). In neuroWalknet, as the main difference, the control circuits as such have been included into these neural circuits, one for each joint. Still, we can identify to a certain degree how the different paths in these circuits elicit swing and stance movement as they drive the corresponding motor neurons. One crucial basic property of these motivation unit networks concerns the structure that allows for switching between different contexts (Fig. 1 shows a part of a single leg controller, for full structure see (Schilling & Cruse, 2020), their Fig. 2) whereby various sensory motor networks (or “sensory-motor primitives”) can be combined in different ways and on different levels.

**Fig. 1.**
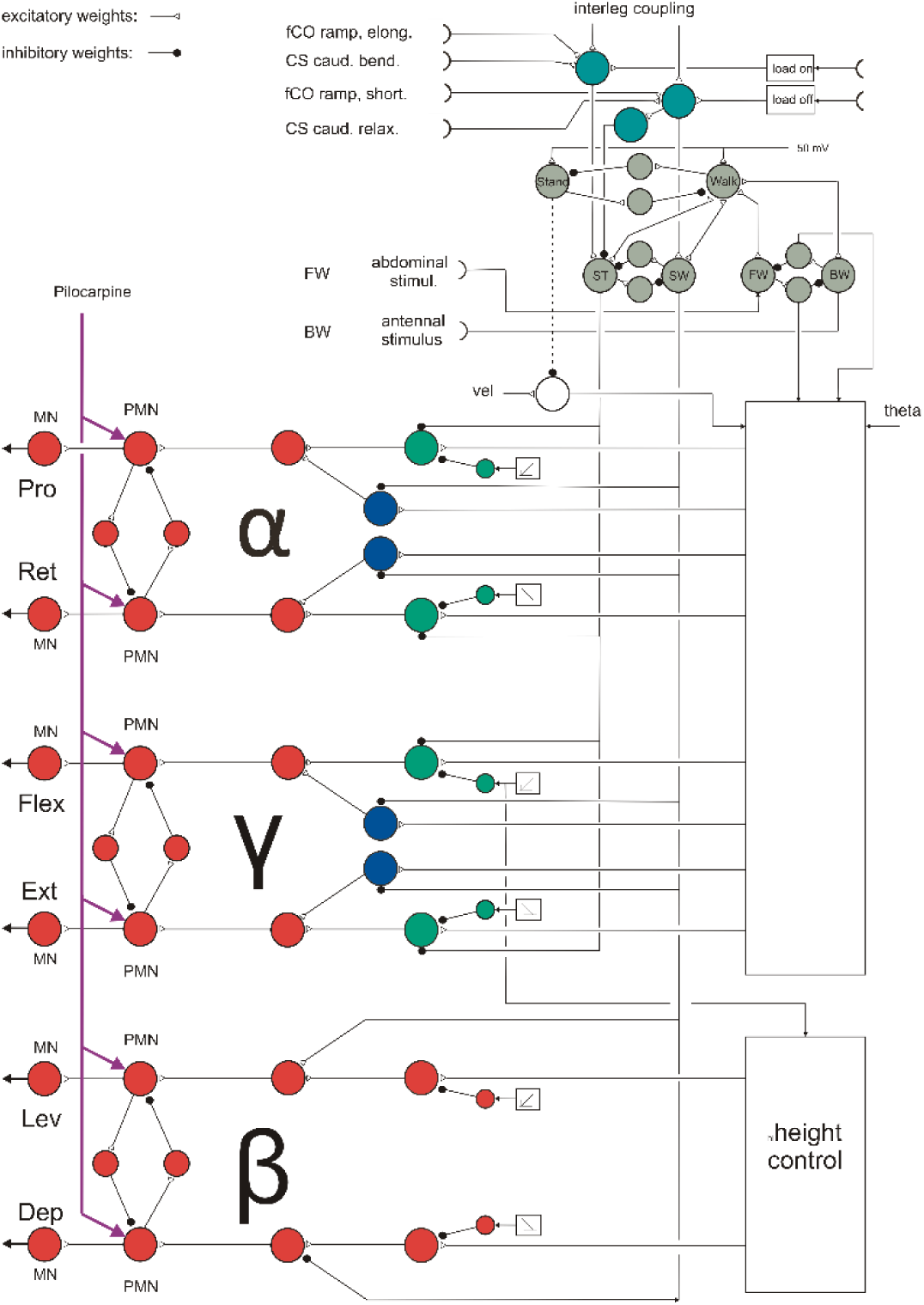
shows a section of neuroWalknet circuitry including the four units representing the bistable monopole controlling forward-backward walking (FW, BW, stimulated by sensors representing the tactile stimulation at the abdomen at the back or antennae at the front which induce walking in the opposite direction, i.e., forward walking or backward walking respectively), and another bistable monopole controlling swing-stance (SW, ST) stimulated by load sensor or position sensor (load on, load off). Unit Stance can further be stimulated by two specific sensors, fCO ramp (stimulus elong.), and CS, caudal bending. Correspondingly, unit Swing can be stimulated by fCO (ramp, short.) and CS (caudal, relax.). Unit Walk is recurrently coupled to units Stance, Swing, FW, and BW. For the third bistable monopole Stand-Walk see text.

**Fig. 2.**
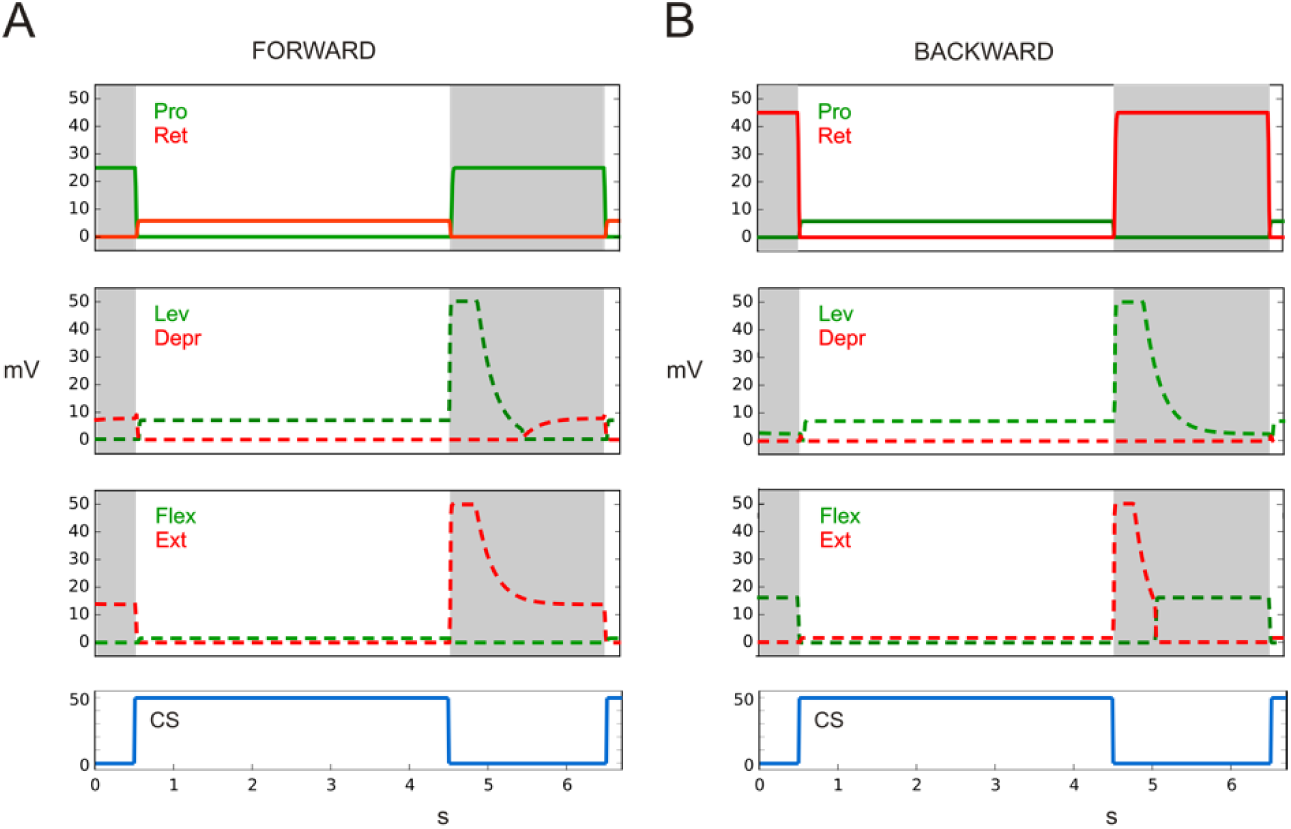
CS stimulus drives (via caudal bending of femur) protractor or retractor as observed in (Akay et al., 2007). Upper lines (alpha joint, or TC joint): Protractor (green), Retractor (red), middle lines (beta joint, or CT joint): Levator (green), Depressor (red), lower lines (gamma joint, or FT joint): Flexor (green), Extensor (red). Below: activation of CS stimulus (blue). A) state: Forward walking, B) state: Backward walking. Abscissa: time (s), ordinate (mV). Dashed lines: simulation data for which no biological results are given.

The “upper” pair controls the states “Stand” or “Walk”, corresponding to passive state or active state (e.g. (Bässler, 1988; Ekeberg et al., 2004)), the two lower pairs control the states “Stance” and “Swing” (Fig. 1, grey units, ST, SW) and “Forward” walking or “Backward” walking (Fig. 1, grey units, FW, BW). All these pairs represent bistable monopoles, which means that only one unit of each pair can be activated at a given moment in time. Using these three pairs of motivation units corresponds to an earlier version of Walknet that was however missing the details of the neural structure for control of the individual joints (Schilling, Hoinville, et al., 2013; Schilling, Paskarbeit, et al., 2013). Note that in the current network (Fig. 1)—as well as in the earlier version ((Schilling & Cruse, 2020), their Fig. 2)—sensory feedback from antennal and abdominal sensors could stimulate unit FW or BW and unit Walk, which in turn activates units Stance or Swing. This is due to the recurrent connectivity of the motivation unit networks.

In neuroWalknet (Schilling & Cruse, 2020), to keep the structure simple, instead of using the pair Walk – Stand we used a single unit (“Leg”), which was possible because we did not address the case “Stand” in these simulations. In the current extension, we used this pair (Walk – Stand) explicitly because these two states are addressed in many of the biological studies used here (cases concerning state “Stand” are addressed in the Supplement). These units provide a form of high-level context.

As illustrated in Fig. 1, activation of unit Stand sets the global velocity to zero (dashed line: inhibitory connection from unit Stand to unit velocity), but other networks not yet implemented here may be activated in the animal. As illustrative examples for state Walk, the retractor muscle (Fig. 1, lower left) may be activated by activation of units Forward and Stance, or by activation of units Backward and Swing. This circuitry is partly hidden in the box (Fig. 1, lower-right), details of which can be found in Schilling and Cruse ((Schilling & Cruse, 2020), their Fig. 2). Green units represent swing activation (upper branch Forward, lower branch Backward). Blue units (upper and lower branches) are required for stance and both forward walking or backward walking depending on actual leg position (and on walking direction when negotiating curves). Red units guide the different channels to the motor output.

Note that the simulated neurons for motor output (e.g., Fig. 1, red units) do not represent the output of individual biological neurons, nor their dynamic properties. Rather, we deal with simplified structures, which means that a comparison between biological data and simulated results is only possible on a qualitative level.

### New sensory inputs

For sensory input to allow for activating states Swing or Stance, in neuroWalknet (Schilling & Cruse, 2020) a minimum solution had been introduced. Touchdown of the leg tip (Fig. 1, load on) activates unit Stance, load off activates unit Swing and inhibits Stance (Fig. 1, three turquoise units, input upper right). These units are further influenced by coordination signals that depend on leg position of the sender leg. In experiments where information from other legs is cut off, these coordination signals are not active.

To enable the network with more specific sensory signals as used in the experiments addressed in the current study, parallel to the “load on” input and “load off” input as applied in neuroWalknet ((Schilling & Cruse, 2020), Fig. 2), two other specific sensory inputs have been introduced. Concerning activation via the CS, a specific input is now introduced that is assumed to be stimulated when the fixed femur is being bent or relaxed ((Akay et al., 2004, 2007; Schmitz, 1993; Schmitz & Stein, 2000), Fig. 1, upper left). Another input to the same units is assumed to be activated while the fCO receives a ramp-wise elongation or shortening of the fCO (e.g. (Bässler, 1976, 1986, 1988; Bässler & Büschges, 1990; Hellekes et al., 2012)). For further details see Methods and (Schilling & Cruse, 2020).

### Variable input representing the effect of Pilocarpine

Another similar, but not identical network is addressed by the four units termed premotor neurons (PMN) in neuroWalknet (Fig. 1, lower left side, and (Schilling & Cruse, 2020), their Fig. 2). It represents a soft winner-take-all net which means that the stronger activated unit suppresses the weaker one by lateral inhibition to minimize co-contraction of both antagonistic branches which is, in particular, relevant when the controller switches between branches. This effect is strengthened by using inhibitory units with phasic properties. As a result, if pilocarpine is given to both units, the network may show cyclic activation. As shown in the section on detailed Methods, various input values representing the activity of pilocarpine have been tested.

Figs 2 – 8 data show simulation results. Sensory inputs are depicted blue. Motor output activation are depicted in red or green color. Continuous lines and dotted lines show motor output that simulates data provided by the biological studies. Dashed lines show motor output that, to our knowledge, has not yet been recorded by experimental studies and may therefore be used as test set in later biological experiments. In figures 7 and 8 the same colors, now in black frames, indicate—for the corresponding branches—input via the pilocarpine activation (for details see Methods).

**Fig. 7.**
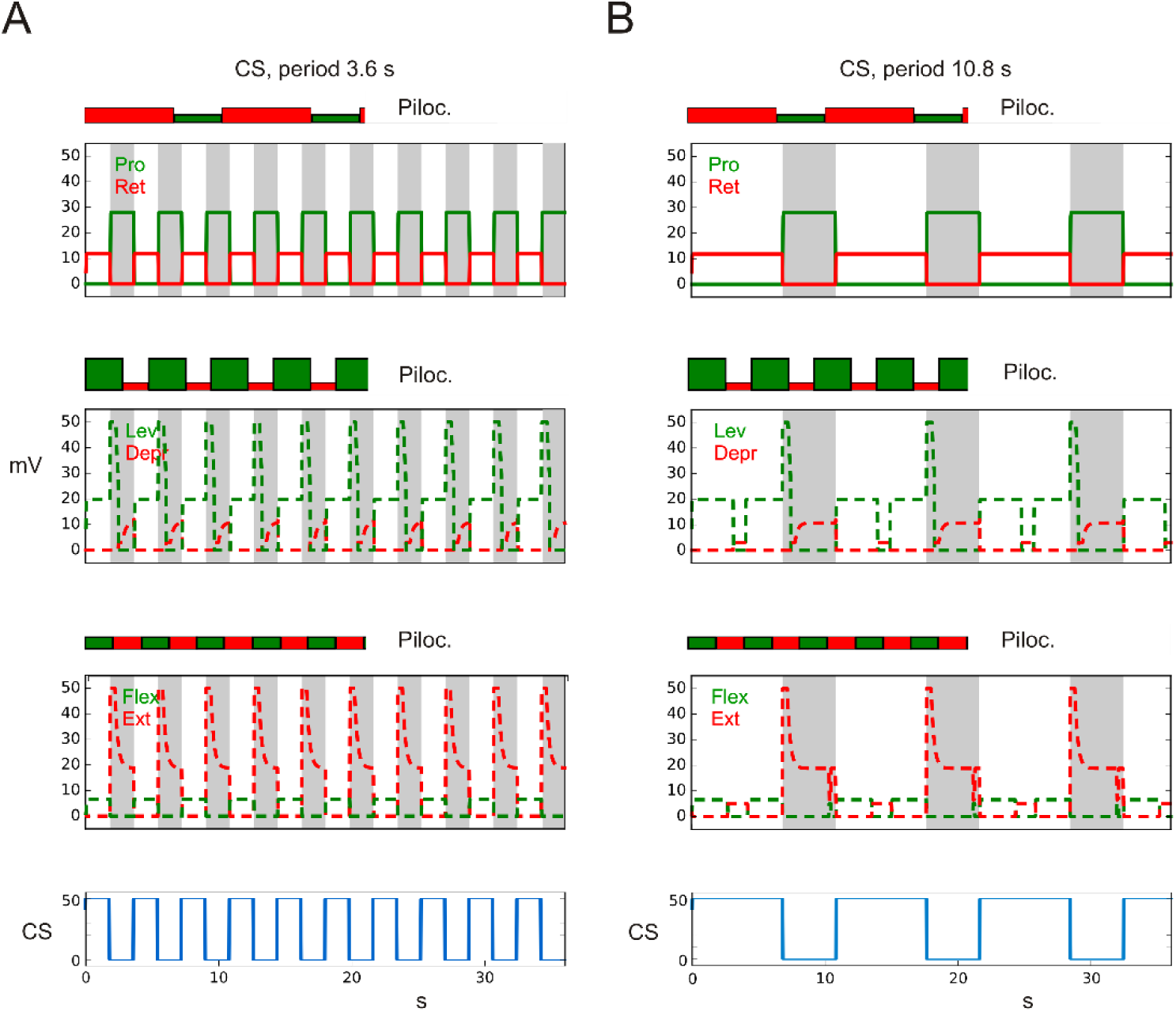
CS stimulus drives (via caudal bending of femur) protractor or retractor as observed in Akay et al. (Akay et al., 2007). Upper lines (alpha joint, or TC joint): Protractor (green), Retractor (red), middle lines (beta joint, of CT joint): Levator (green), Depressor (red), lower lines (gamma joint, or FT joint): Flexor (green), Extensor (red). For comparison, cycles that are driven only by pilocarpine are plotted for each joint on top of the CS-driven results (same scale, periods: alpha joint: 10.2 s, beta joint 4.6 s, gamma joint: 4.2 s) and marked by the same colors but with filled rectangles and ‘piloc.’. Below: activation of CS stimulus (blue), state: forward walking, A) Sensory input, period: 3.6 s, B) Sensory input, period: 10.8 s. Abscissa: time (s), ordinate (mV). Dashed lines: simulation data for which no biological results are given.

**Fig. 8.**
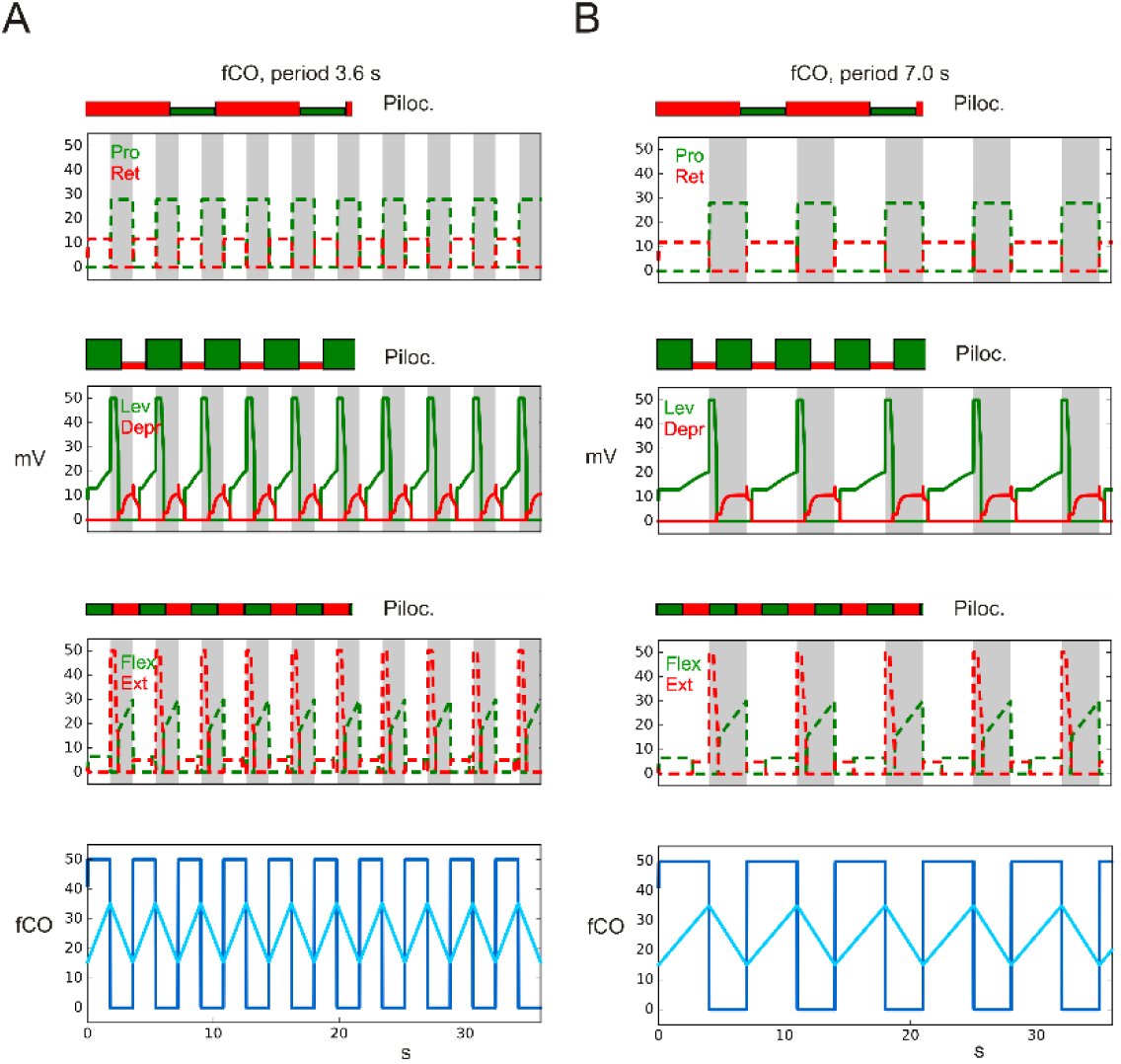
fCO stimulus (elongation, shown in bottom row as light blue line) drives Levator or Depressor as observed in Hess and Büschges (1999). Upper lines (alpha joint, or CT joint): Protractor (green), Retractor (red), middle lines (beta joint, or TC joint): Levator (green), Depressor (red), lower lines (gamma joint, or FT joint): Flexor (green), Extensor (red). For comparison, cycles driven by pilocarpine only are plotted for each joint on top of the fCO-driven results (same scale). The former are marked by the same colors but with filled rectangles and ‘piloc’. Below: activation of fCO stimuli (blue), state: forward walking, A) Sensory input, period: 3.6 s, B) Sensory input, period: 7.0 s. Abscissa: time (s), ordinate (mV). Dashed lines: simulation data for which no biological results are given. Gamma flexion corresponds to fCO apodeme elongation, fCOpos: position input to Levator-Depressor joint (gamma joint), light blue line indicates fCO apodeme position, fCO: (velocity) input to unit swing or unit stance (dark blue line).

Although in the various biological experiments different legs have been studied, here we use front legs for simulation of each experiment and will comment in the Discussion if experiments with other legs show different results.

## 3 Results

In the following, we will briefly recall the four different groups of experiments mentioned in the Introduction (see also Table 1) and then show for each case how neuroWalknet will react to the corresponding experimental situations. We will not only show the simulation results for the motor activations as recorded in the biological experiments, but will show the data of all three simulated joints. These additional results (shown by dashed lines) can therefore be considered predictions for possible future experiments.

In all experiments addressed here, biological and our simulation experiments represent open loop experiments, i.e., the sensory input to CS or fCO is controlled by the experimenter. For results of closed loop experiments (e.g., free walking as forward or backward or negotiating curves) see (Schilling & Cruse, 2020), their figures 3, 4, 7, 8, and corresponding videos.

**Fig. 3.**
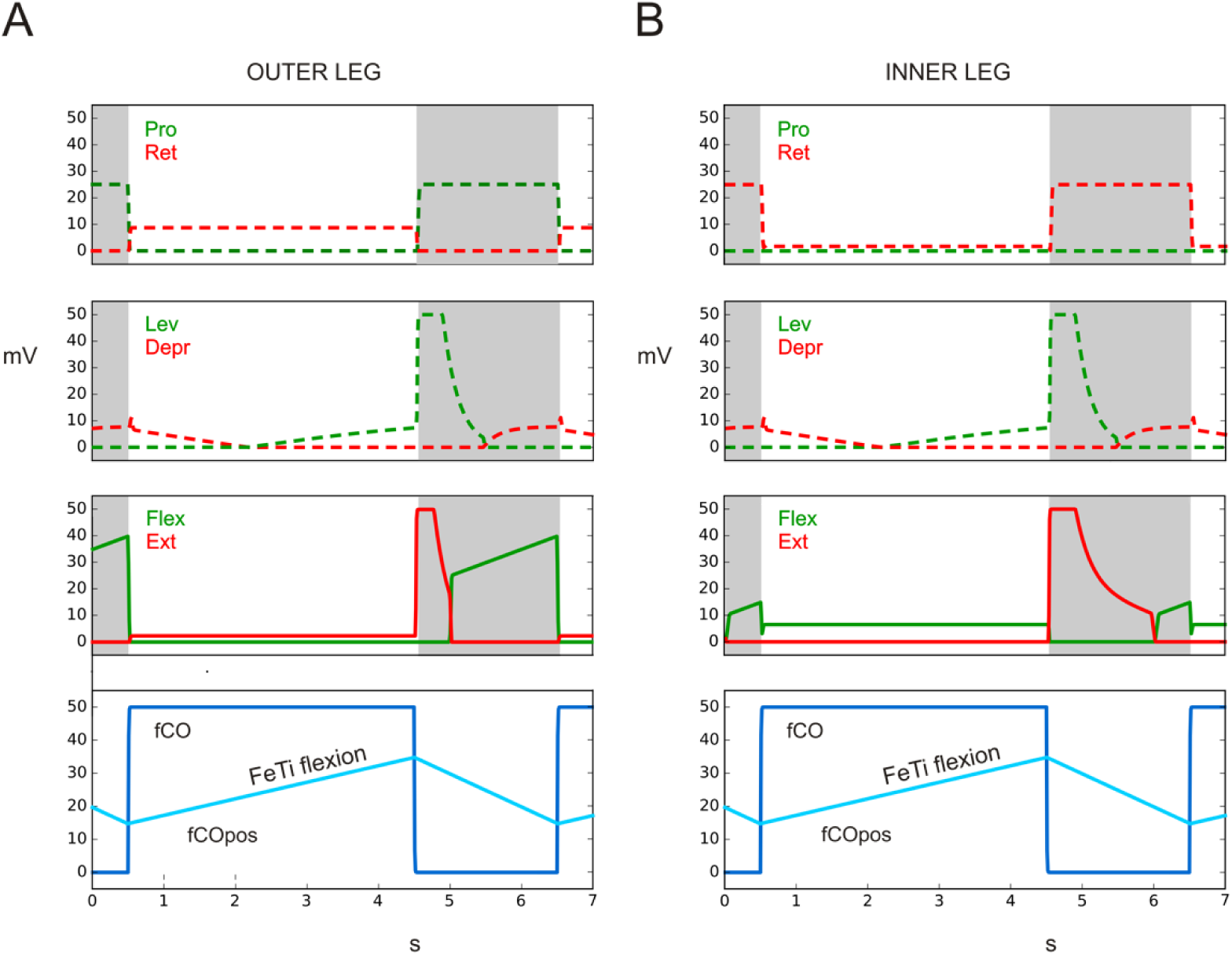
Negotiation of curves (turn to the right). fCO stimulus (elongation) drives flexor or extensor as observed in (Hellekes et al., 2012), but here front legs are shown. Upper lines (alpha joint, or CT joint): Protractor (green), Retractor (red), middle lines (beta joint, or CT joint): Levator (green), Depressor (red), lower lines (gamma joint, or FT joint): Flexor (green), Extensor (red). Below: activation of fCO stimulus (blue). A) outer (left) front leg, B) inner (right) front leg. Abscissa: time (s), ordinate (mV). Dashed lines: simulation data for which no biological results are given. fCOpos (light blue line) indicates fCO apodeme position, FeTi shows direction of movement. fCO: input to unit swing or unit stance.

**Fig. 4.**
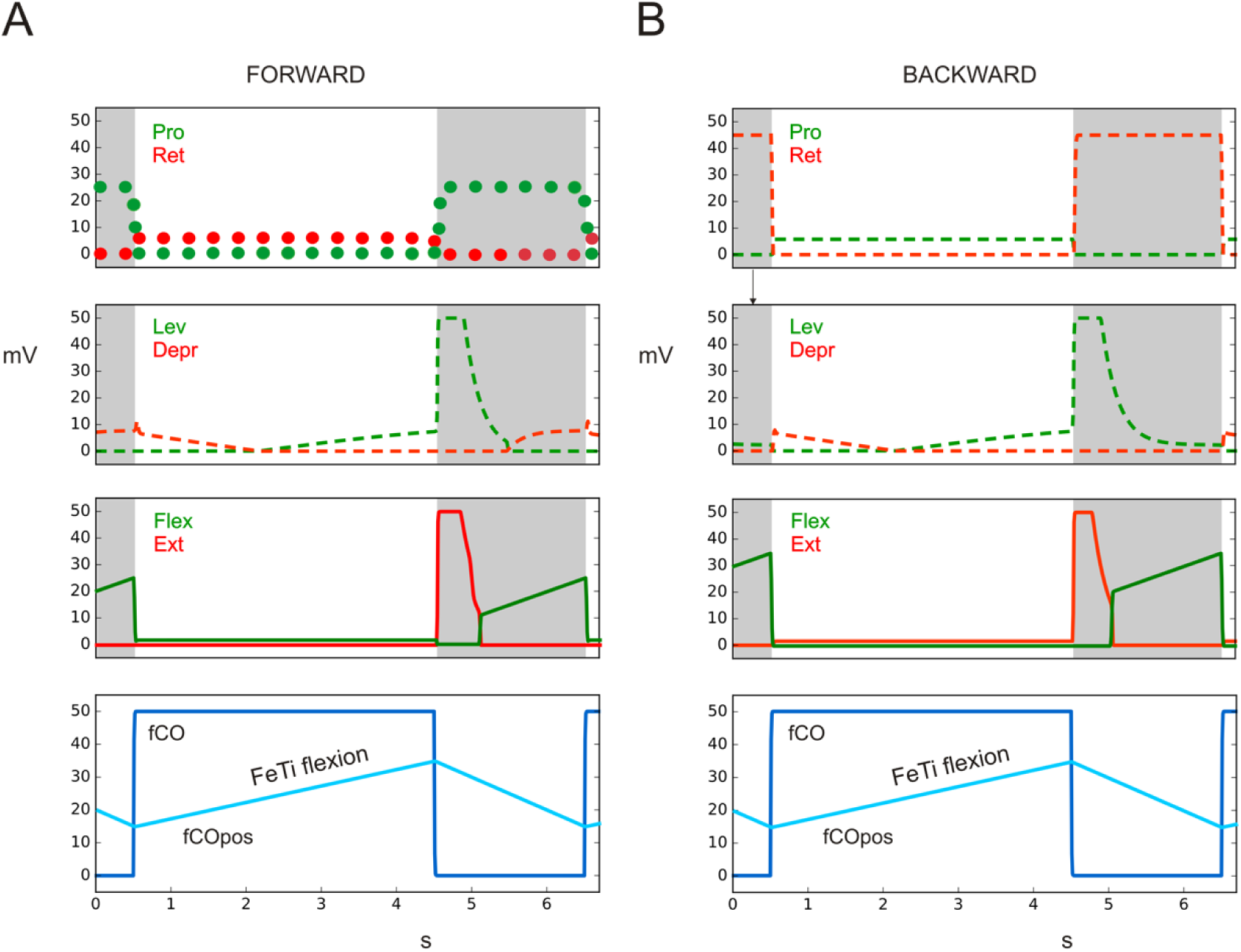
Active reaction. fCO stimulus (elongation) drives flexor or extensor as observed in (Hellekes et al., 2012). Upper lines (alpha joint, or TC joint): Protractor (green), Retractor (red), middle lines (beta joint, or CT joint): Levator (green), Depressor (red), lower lines (gamma joint, or FT joint): Flexor (green), Extensor (red). Below: activation of fCO stimulus (blue). A) state: forward walking. B) state: backward walking. Abscissa: time (s), ordinate (mV). Dashed lines: simulation data for which no biological results are given. Dotted lines: results from (Bässler, 1986). fCOpos (light blue line) indicates fCO apodeme position, FeTi flexion shows direction of movement. fCO: input to unit swing or unit stance.

### 3.1 How stimulation of campaniform sensilla drives retractor and protractor units in the alpha joint depending on context (fw–bw) during active state

<u>Biological experiments:</u> In an interesting paradigm, Schmitz and colleagues (Akay et al., 2004, 2007; Haberkorn et al., 2019; Schmitz, 1993; Schmitz & Stein, 2000) stimulated load sensors—the campaniform sensilla—placed near the beta joint and recorded activation of retractor and protractor of the alpha joint of the middle leg. While the coxa was fixed to the body, and deafferented except for input from the campaniform sensilla, the latter were stimulated by bending the femur caudally or rostrally. In the passive standing animal, caudal bending elicited activation of retractor muscle, whereas rostral bending elicited protractor activation. When the animal was activated by tactile stimulation to reach an active state, the response was in principle the same but showed much stronger activation. (Akay et al., 2007) extended this paradigm by studying not only middle leg and hind leg, but also front legs. In addition, they introduced context changes by using tactile stimulation of abdomen or antennae to elicit forward or backward walking, respectively, while animals walked on a slippery glass plate or were fixed with legs being removed. The experiments showed that, with a fixed leg stimulated with femoral bending and the other legs being intact or removed, as above, caudal bending elicits retractor activation during forward walking. But when walking direction is changed to backward walking, caudal bending lead to an immediate switch from retraction to protraction thus showing a context dependent reflex reversal ((Akay et al., 2007), their figures 1, 4). When legs are removed, however, hind legs show an inverse behavior which will not be considered here but dealt with in the Discussion.

### Simulation

How does neuroWalknet react in these situations? In the simulations studied here, the leg tested is in active state. If the leg is not stimulated by bending, i.e., if there is no stimulation by load sensors, this leg is assumed to be in state Swing. As soon as the leg is bend, this stimulation triggers the state Stance (Fig. 1) and thereby inhibits unit Swing.

As a result, the simulation with neuroWalknet, when stimulated by caudal bending (Fig. 2), indeed shows that when the leg is in state forward, the retractor is activated (Fig. 2a, red, filled lines), while when in state backward, by the same stimulus now the protractor is activated (Fig. 2b, green, filled lines), as observed in the biological experiments ((Akay et al., 2007), their Fig. 4). In both cases, relaxation of CS stimulation leads to activation of the antagonistic muscle.

In both simulation experiments, the switch from state Forward to state Backward and back immediately influences the motor output as observed in the biological experiments ((Akay et al., 2007), their Fig. 7). Here we show only simulation of a front leg, as in principle the effect of context dependency (walking direction, Forward or Backward) can be seen in all three legs (for hind leg, see Discussion).

### 3.2 How stimulation of fCO drives flexor and extensor units (and protractor and retractor units) depending on contexts (negotiation of curves, forward–backward walking) in active state

#### Biological Experiment

The **Resistance Reflex**, or a negative feedback response, may be observed, when in a quiet animal—i.e., legs are not actively moved as, for example, no tactile body stimulation is given—the gamma joint (FT joint) is flexed or extended by the experimenter or the fCO apodeme (or tendon) is experimentally elongated or shortened, respectively. Elongation of fCO (i.e., flexion of the gamma joint) leads to activation of extensor muscle, i.e., to extension of the gamma joint, whereas shortening the fCO apodeme (i.e., extension of the gamma joint) leads to activation of the flexor muscle.

The second intensively studied aspect concerns the so-called **Active Reaction**. In this paradigm (Bässler, 1976, 1986, 1988; Bässler & Koch, 1989; Nothof & Bässler, 1990) the animal was fixed and legs had no ground contact. Usually, one leg was restrained and the fCO could be stimulated by a rampwise elongation of the fCO apodeme. Following tactile stimulation of the abdomen, which elicits forward walking, elongation of fCO apodeme elicits flexor activation which is in direct contrast to the reaction described above for the resistance reflexes that lead to extensor activation.

If a given elongation has been reached, flexor activation is switched off being replaced by extensor activation. The former part has been called Active Reaction I, the second Active Reaction II, whereby the latter has been interpreted as starting a swing movement.

The type of behavior observed in these studies depends on the context, on a higher level being passive state or active state. At a lower level, when in active state, during negotiation of curves, the context may depend on the leg walking on the inner side or the outer side of the curve, or, may depend on state forward or backward. Our simulations only consider distinctions concerning active state, i.e. curve walking and forward – backward walking.

Hellekes et al. (Hellekes et al., 2012) continued these studies further. They turned to experiments in the active animal, and our focus will be on these experiments. They investigated Active Reaction with most legs being intact and walking on a slippery plane, while only one leg was fixed so that the fCO of this leg could be stimulated experimentally. Forward walking and backward walking as well as curve walking have been studied. Here, we will focus on experiments that can be observed in the active state. Properties of behavior belonging to the passive state will be addressed in the Discussion. First, we will consider negotiation of curves as a context which represents quite a complex behavior and where embodiment plays an important role (for free walking see (Schilling & Cruse, 2020), their Fig. 6, and video, and (Dürr & Ebeling, 2005; Rosano & Webb, 2007)). In the second part we will focus on straight walking and compare forward walking and backward walking.

**Fig. 6.**
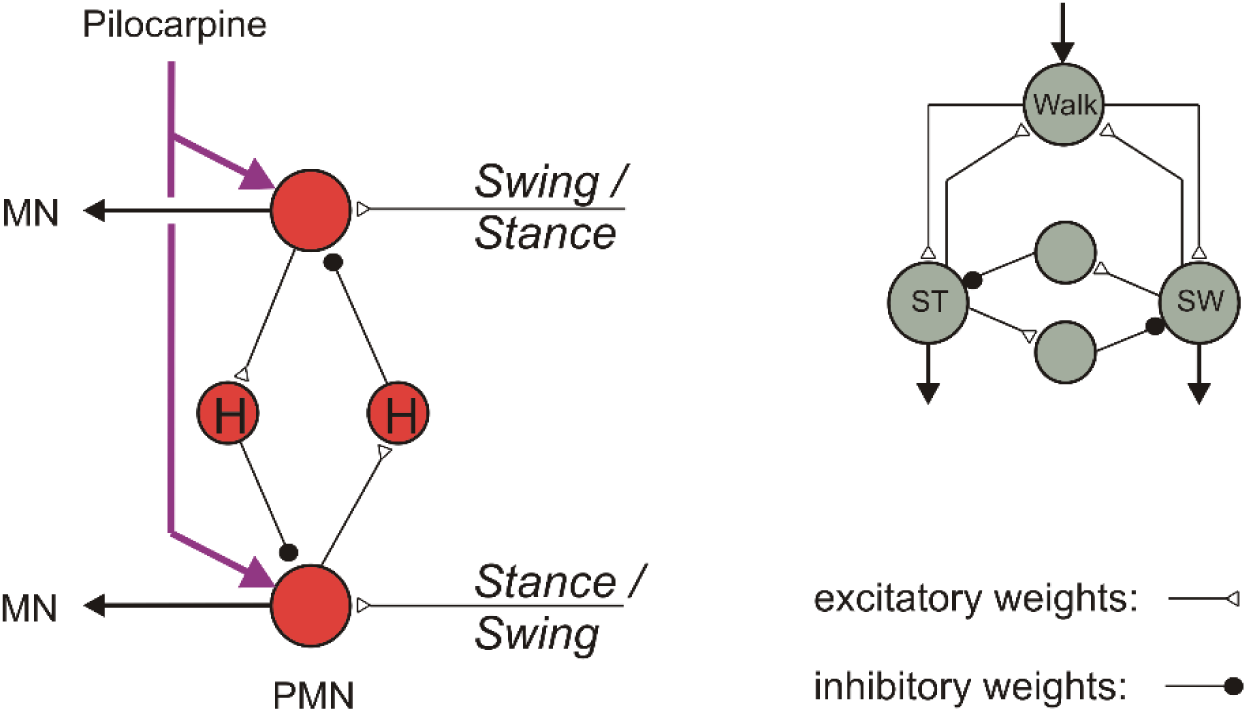
Two similar but not identical networks that are elements of units shown in Fig. 1. Left: Four units (red) form a PMN network. Both inhibitory units are equipped with a high-pass filter (H). The purple arrows represent influence of pilocarpine. Right: four units (grey), two of which are recurrently connected with unit Walk via constant input. Both inhibitory units do not have high-pass properties, but only low-pass filter properties as is the case for all the neurons in neuroWalknet.

### 3.2.1 Negotiating curves – context dependent influence on different legs

#### Biological Experiments

In the biological experiments (Hellekes et al., 2012), animals have been stimulated by visual input to negotiate curves with five legs walking on slippery ground and with the left middle leg being prepared to be tested for Active Reaction. When fCO of the inner leg was experimentally flexed (i.e., fCO apodeme was elongated), an Active Reaction, i.e., a flexor activation was observed. Thus, the Active Reaction observed supports the movement which is expected during stance because flexor action is also observed in the intact inner leg during curve walking. For the outer middle leg the situation is different ((Hellekes et al., 2012), their Fig. 7).

During stance typically the gamma joint is either about constant or, depending on curve radius, produces an extension. When in the corresponding experiments the fCO was elongated, Active Reaction could not be observed in the biological experiment ((Hellekes et al., 2012), their Fig. 7), as the data did neither support a clear flexor activation nor a clear extensor activation. This result might, however, be expected functionally, because stance in the outer middle leg does not anymore perform a flexor movement, rather, depending on curve radius, an extension.

#### Simulation

For the **simulations of curve walking,** as in the other cases studied here, front legs have been used. Specifically, for the curve radius used here this has the advantage that front legs show a qualitatively similar but stronger effect during curve walking than do middle legs. As an example, for a walking trajectory turning to the right, we apply a turning angle of theta = 75 deg as used by ((Schilling & Cruse, 2020), their Fig. 6), the morphological arrangement of which is based on data from (Dürr & Ebeling, 2005) and approximately corresponds to the behavior observed by (Hellekes et al., 2012).

In the experiment shown in Fig. 3, unit Walk is activated. As is the case for all simulations, Fig. 3 shows the corresponding motor neuron activations for all three joints. Fig. 3A provides the corresponding results for the outer leg. When the fCO is elongated, corresponding to the flexion of the gamma joint, a weak extension is observed (red filled lines) For the inner leg a corresponding simulation is given in Fig. 3B. Here elongation of the fCO (i.e. the gamma joint is flexed) leads to an activation of the flexor muscle (green filled lines). Qualitatively, both results agree with the findings of ((Hellekes et al., 2012), their Fig. 7). These results support the idea that the structure of neuroWalknet combined with the physical properties of the body may allow for the control of insect-like walking behavior as studied here. Specifically dedicated neuronal elements for Active Reaction may not be required as a separate structure. Rather, this behavior may be considered to emerge out of the complete system. In other words, a specific system for switching on or off an Active Reaction appears not necessary for curve walking.

### 3.2.2 Straight walking depending on context (fw–bw)

#### Biological experiments

In the following, we concern how the gamma joint may contribute to forward walking and backward walking where, as above (Fig. 2) these two states are activated by tactile stimulation of abdomen or antennae, respectively. As shown by (Hellekes et al., 2012) (their Fig. 5), when in state forward walking stimulation of the fCO of a front leg by elongation elicits an Active Reaction, i.e., activation of the flexor. This behavior would support a stance movement in the case of an intact walking leg. When, however, the animal is stimulated to walk backward, during experimental elongation of fCO apodeme no Active Reaction, i.e., no flexor movement has been observed. Rather, the extensor is activated instead, which appears functionally sensible because extension would support stance during backward walking (for results concerning the hind leg, see (Nothof & Bässler, 1990), Discussion).

**Fig. 5.**
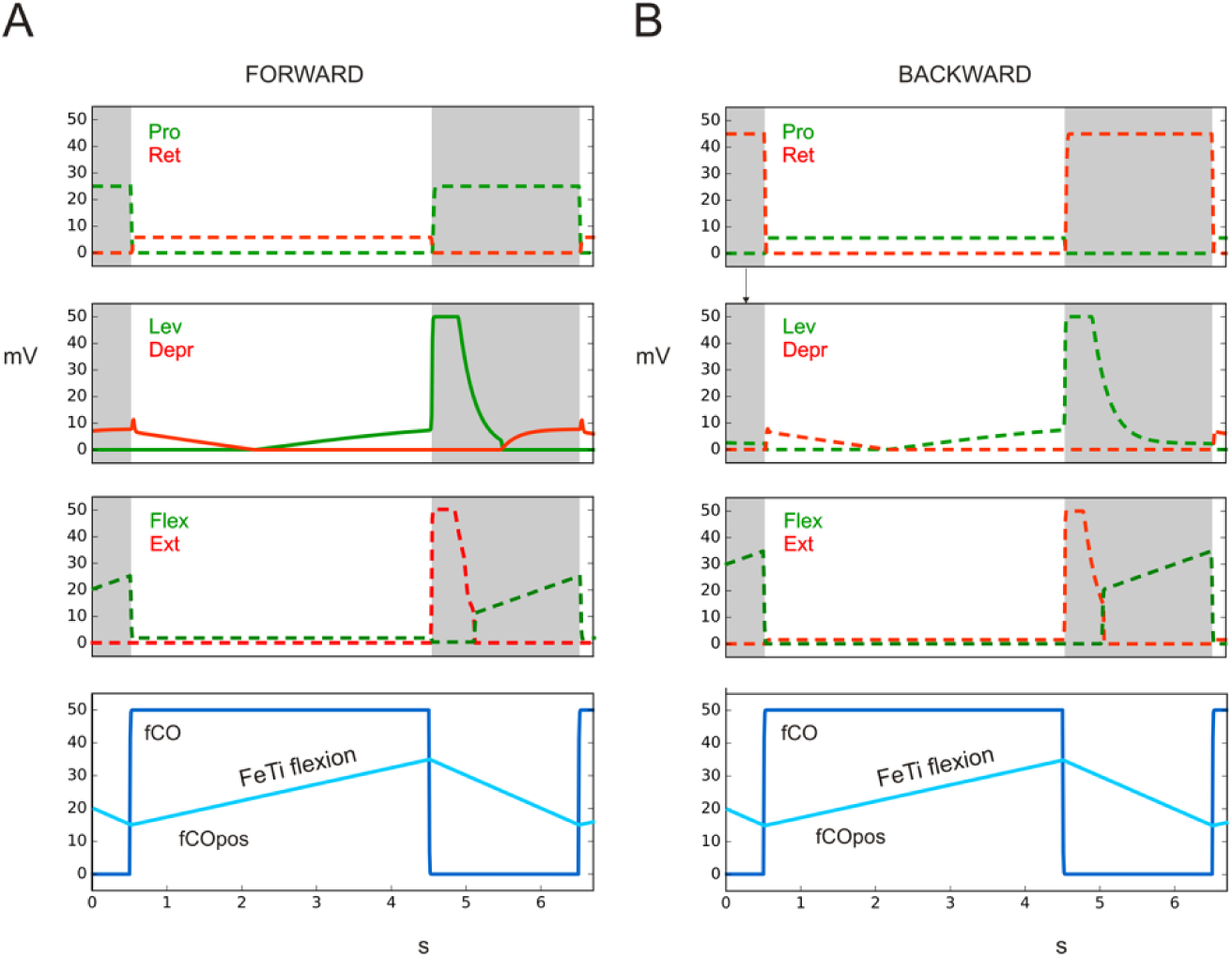
fCO stimulus (elongation, shown below as dark blue line) drives Levator or Depressor as observed in Hess and Büschges (1999). Upper lines (alpha joint, or CT joint): Protractor (green), Retractor (red), middle lines (beta joint, or CT joint): Levator (green), Depressor (red), lower lines (gamma joint, or FT joint): Flexor (green), Extensor (red). Below: activation of fCO stimulus (blue). A) state forward walking. B) state backward walking. Abscissa: time (s), ordinate (mV). Dashed lines: simulation data for which no biological results are given. fCOpos (light blue line) indicates fCO apodeme position, FeTi flexion shows direction of movement, which, via the height controller, activates the Levator-Depressor system (beta joint). fCO: input to unit swing or unit stance.

#### Simulation

How would neuroWalknet react in this situation? When state forward walking is activated and the fCO of a front leg is stimulated by elongation, then—due to the structure of neuroWalknet—the flexor is activated (Fig. 4A, green filled lines). Correspondingly, when state backward walking is activated, the extensor is activated (Fig. 4B., red filled lines). Both results are in qualitative agreement with results of Hellekes et al. (2012, their Fig. 5). The effect is relatively weak due to the smaller angular range used during normal walking.

Interestingly, (Bässler, 1986) performed a detailed study with the front leg of *Cuniculina* and recorded not only the gamma joint as simulated by (Hellekes et al., 2012), but also the alpha joint, i.e., activation of retractor and protractor. He found that in forward walking flexor activity was accompanied by increase in retractor activity which, too, agrees with the neuroWalknet simulation results when walking forward (Fig. 4A, red dotted lines). Our simulation predicts that changing walking direction to backward walking will activate protractor during stance (Fig. 4B, green dashed lines).

Taken together, all the experimental results reported by (Hellekes et al., 2012) and mentioned here (including data from (Bässler, 1986)) are in qualitative agreement with the simulated behavior produced by neuroWalknet. In other words, what has been described as Active Reaction could now be explained as a property emerging from a general, decentralized structure as given in neuroWalknet (see Discussion).

### 3.3 fCO stimulation drives levator units and depressor units, but the active state does not elicit reflex reversal

#### Biological experiments

After we have seen cases of reflex reversal in the alpha and gamma joint depending on the context, in the third group of experiments we turn towards the beta joint. In one series of experiments the influence of stimulating the fCO onto the levator-depressor system (beta joint) has been investigated (Hess & Büschges, 1999). In these experiments front legs and hind legs were fixed. The stimulated middle leg underwent basically the same preparation as above (Hellekes et al., 2012): elongation of fCO, and tactile stimulation of the body to trigger active state. But now activation of levator motor neurons and depressor motor neurons of the beta joint have been recorded. Different to recordings from both systems treated above—the protractor – retractor system or the flexor – extensor system—, in the beta joint no reflex reversal could be observed, i.e. there was no Active Reaction when the animal was activated by tactile stimulation of the body. Rather, elongation of the fCO shows an increasing activation of levator in both states, active or passive. There were only some minor differences: Latencies were larger and more variable in the active state. Activation of levator muscles was somewhat stronger in the active state than in the passive state, but did not show a dependency on several stimulus variables, e.g. position, or velocity.

In a later study, Bucher et al. (Bucher et al., 2003) provided more quantitative details in the active state. In particular, they found a nonlinear dependency between fCO position and corresponding beta angle values, which was complicated by nonlinear dynamic properties.

#### Simulation

As in the earlier simulations, we will only focus on active state. In neuroWalknet, the beta controller contributes to height control, the function of which is to keep body-ground distance approximately constant independent of tibia position. The beta controller only refers to positional input, as it was only tasked to control body height. Therefore, during stance, positional input from gamma joint influences the beta angle (as shown in (Schilling & Cruse, 2020), Fig. S1) and elongation of fCO (flexion) activates the levator as long as—in the case of neuroWalknet—the gamma remains in a range of > 60 deg. Therefore, different to observations for the alpha and gamma joints, neuroWalknet is not expected to show a reflex reversal in the beta joint.

The corresponding simulation results for forward walking are shown in Fig. 5A (red/green filled lines) and qualitatively agree with data of (Hess & Büschges, 1999) (their Fig. 1). Note that neuroWalknet is characterized by an antagonistic neuronal structure. Decrease of depressor (red lines) follows an increase of levator (green lines) and both agonists move the output in the same direction.

In these biological experiments, only forward walking has been studied, but following the architecture of neuroWalknet, a different influence of backward walking is not expected, as the beta controller in neuroWalknet does not receive input from the forward-backward network ((Schilling & Cruse, 2020), their Fig. 2). Thus, Fig. 5B (backward walking) shows the same results for the beta joint (red/green filled lines).

Nonlinearities as described by Bucher et al. (Bucher et al., 2003) have not been implemented as these are not required for the small angular range observed during walking (Schilling & Cruse, 2020).

As mentioned in (Hess & Büschges, 1999) (their Fig. 2), latencies are more variable in the active state then in the passive state. This result might be explained if the neuronal structure follows the principle structure of neuroWalknet. In neuroWalknet, when representing the active state, both the height controller as well as activation via units Swing or Stance can influence the input of levator and depressor. However, in the passive state only part of neuroWalknet, the height controller, may by activated. Therefore, the active state may lead to a higher variability, as observed by (Hess & Büschges, 1999).

### 3.4 How sensory input (CS, fCO) influence artificially induced rhythmic oscillation

In the fourth group of experiments, we will test how oscillations that have been experimentally induced (Akay et al., 2007; Hess & Büschges, 1999) can be recreated by neuroWalknet. These and earlier studies have often been seen as an argument for requiring an explicit central pattern generator structure. Such a structure is not fully realized in neuroWalknet. Therefore, the fourth group of experiment is as well a test if such oscillations do exist in the biological system as a necessary element for control of walking or if the present structures in neuroWalknet that do not operate with CPGs, i.e. not treated with pilocarpine, are sufficient to explain such a behavior.

#### Biological experiment

In the following experiments, legs did not only receive tactile stimulation of abdomen or antennae and CS or fCO stimulation as in the above experiments, but were in addition treated with pilocarpine. Pilocarpine induces an overall activation that shows as internal rhythmic movements of low frequency. In the classical experimental studies (see e.g. (Büschges et al., 1995)) sensory input and motor output as well as connectives have been cut. In contrast, in the current experiments (Akay et al., 2007; Hess & Büschges, 1999) sensory stimulation is made possible by leaving the neuronal connections to CS or to fCO intact in order to permit additional external rhythmic or arhythmic stimulation.

Importantly, in both of these biological experiments—addressed here and as shown in Fig. 7 and Fig. 8—the sensory stimulation was rhythmic and characterized by periods shorter than that of the periods produced by the pilocarpine driven oscillations.

#### Realization of experimental influence in simulation

How is the experimental effect provided by pilocarpine introduced into neuroWalknet? First of, we consider the neural structure that is present in neuroWalknet. In neuroWalknet, a Winner-take-all like structure of competing units can be found in multiple instances. In addition to the competing units, two additional units are used to organize the activations of the other two units through mutual inhibition. These structures are used in two different contexts. First, one of these structures is present for each joint in each leg network and this has been called the premotor neuron net (PMN), controlling the input to the motor neurons (MN) of its joint. Such a PMN net (three in each leg) contains these four neuronal units each, two excitatory, and two inhibitory ones (units in red, Fig. 1, Fig. 6). Secondly, such a network—that also consists of four units—is employed as part of the motivation unit network, for example, the net that controls the switch between Swing and Stance (SW – ST, grey units in Fig. 1, Fig. 6), which is called a bistable monopol. Importantly, there are two differences in the incoming connectivity of these networks. In the motivation network on the leg level, as in the SW – ST net, both excitatory units receive a stable constant input, which is not the case for the PMN network. Instead, the inhibitory units of the PMN net contain a high-pass filter, i.e. phasic properties, each (Fig. 6, marked H), which in turn is not the case for the SW – ST motivation unit net.

While these networks represent two different types, they have in common the property that neither of these networks is able to show oscillations on its own (for more details see (Schilling & Cruse, 2020)), because this structure misses the constant excitatory input as used for the motivation unit structure. This changes for the PMN net, if its excitatory units are activated by additional, constant positive excitation elicited by pilocarpine being provided to the ganglion. Depending on the activation strength driving both units, the PMN net may oscillate in different periods and different phase relations. These input channels are shown as purple arrows (Fig. 1, Fig. 6) and represent, in the simulation, the strength of the pilocarpine influence to the PMN unit (for details see Methods, Table 2), i.e., an externally and purely experimentally driven additional input. Note that in the biological experiments this procedure would require surgical treatment.

**Table 2:** Setting of input parameter and the corresponding periods for the fourth experiment (pilocarpine application).

|  | Protr. | Retr. | Period | Lev. | Depr. | Period | Flex. | Ext. | Period |
| --- | --- | --- | --- | --- | --- | --- | --- | --- | --- |
| BSB (s) | 2 | 4-7 | 6-9 | 3.7 | 2.7 | 6 |  |  | 0-4 |
| Simulation of current Pilocarpine activation |  |  |  |  |  |  |  |  |  |
| input (mV) | 3.0 | 6.0 |  | 13.0 | 3.0 |  | 5.0 | 5.0 |  |
| output Pilo(s) | 3.7 | 6.7 | 10.2 | 2.8 | 1.8 | 4.6 | 2.1 | 2.1 | 4.2 |
| Simulation of Pilocarpine activation used by Schilling and Cruse (2020) |  |  |  |  |  |  |  |  |  |
| input (mV) | 40.0 | 25.0 |  | 40.0 | 25.0 |  | 50.0 | 50.0 |  |
| output Pilo(s) | 6.4 | 5.4 | 11.8 | 2.0 | 5.6 | 7.6 | 2.0 | 2.1 | 4.1 |

#### Details Biological Experiment 1

We start with experiments of (Akay et al., 2007) showing that after pilocarpine is applied, adding rhythmic (or arhythmic) caudal bending of femur as described earlier (Sect. 3.1) can entrain the motor output signals during both forward walking or backward walking. In the biological experiment ((Akay et al., 2007), their Fig. 1E, CS period about 4 s, pilocarpine driven period about 3 s) for a forward walking middle leg, caudal bending has driven retractor activation. In this case, indeed the CS signal dominates, i.e., resets the pilocarpine induced oscillation if the CS signal has a somewhat shorter period (i.e., higher frequency) than the pilocarpine driven cycle.

#### Details Simulation 1

How can this be simulated? In neuroWalknet (Schilling & Cruse, 2020), treatment with pilocarpine is simulated in the following way. In each leg joint, there is an independent pair of antagonistic premotor neurons (PMN), which are coupled via inhibitory units being equipped with phasic properties, or high-pass properties (Fig. 1, Fig. 6, PMN). The PMN network, when activated by pilocarpine, elicits independent oscillatory rhythms, one for each joint, that activate the corresponding motor neurons. Different frequencies can be chosen through the variation of input values (Fig. 1, Fig. 6, purple arrows, and Table 2). Note, that if these input values are set to zero, no oscillation will occur. As in the biological experiments, connectives to neighboring ganglia, i.e., rule 1-5, have been switched off.

As a second element—already mentioned above (Fig. 1, turquois units)—, the CS signal activates unit Stance or unit Swing, which activates further units via the antagonistic branches (Fig. 1, green units and blue units, see also (Schilling & Cruse, 2020)) and thereby in turn influence the PMNs, too. To summarize: Both inputs—one produced by the pilocarpine treatment, the other by the sensory signals—are combined in the PMN structure, and the result will then be projected to the motor neurons.

To better understand as to how these two inputs are integrated in the PMN structure, we show three different graphics. First, the motor output provided by the network without using pilocarpine as shown in Fig. 2. Second, Fig. 7 combines two further graphics where, for each joint, the outputs are plotted pairwise on top of each other. The respective upper plot shows the motor output data for each joint when driven by pilocarpine only, i.e., without sensory input (upper graphics, marked by rectangles filled red or green and framed black, and by ‘piloc’). To this end, different periods (selected to be similar to frequencies used by (Schilling & Cruse, 2020)) are used (10.2 s (alpha joint), 4.6 s (beta joint), and 4.2 s (gamma joint)), see Table 2. The corresponding lower graphic, for comparison, shows the result of the situation where both—rhythmic sensory input as well as rhythmic input from the oscillation produced by the pilocarpine stimulation—are superimposed. To gain some variability, we used different periods for the sensory input. Fig. 7A shows a CS period of 3.6 s and Fig. 7B a period of 10.8 s. The output data are shown in colors as used in the earlier figures for Protractor – Retractor, Levator – Depressor, and Flexor – Extensor.

Note that due to the reset effect of the CS input, the pilocarpine driven cycles (above) and the combined data are in-phase only for the first period. As mentioned, in Akay et al. (Akay et al., 2007) and in Hess and Büschges (Hess & Büschges, 1999) rhythmic sensory stimulation has been chosen using shorter periods than those of the pilocarpine driven oscillations. In the simulations, we will however also test cases where pilocarpine driven periods are shorter than sensory driven periods. Such results may then be used for further biological studies as a test. Due to the selection chosen, each of these two experiments (Fig. 7A, 7B) contains altogether 6 different tests (2 CS cycles x 3 pilocarpine driven cycles). As a result, in all three joints of Fig. 7A and the alpha joint of Fig. 7B, the simulated data agree with the biological data as the CS input dominates all four cases. In Fig. 7A, in all cases the CS period is shorter than that of the pilocarpine induced periods. In Fig. 7B, alpha joint, the CS period is by about 6 % longer than the pilocarpine driven period, but still the CS signal is dominating. In the beta joint and gamma joint of Fig. 7B, the CS cycle only partly dominates, as the CS cycles are slower than the corresponding pilocarpine driven cycles (sensory period: 10.8 s, for the beta joint: pilocarpine period 4.6 s, for the gamma: pilocarpine period 4.2 s. Therefore, the faster pilocarpine cycles dominate within the ongoing CS period, but still, in both cases, the changes of the CS signal nonetheless reset the output values.

<u>Details Biological Experiment 2:</u> As another, second biological experiment, we use the approach of (Hess & Büschges, 1999) already shown above (Fig. 5). As a further expanded experiment, (Hess & Büschges, 1999) treated the middle leg with pilocarpine ((Hess & Büschges, 1999), their Fig. 8). As expected, oscillations were observed in the levator-depressor system (Büschges et al., 1995). Rhythmic or arhythmic fCO stimulation—introduced by elongation or shortening of fCO apodeme— was able to drive activation of levator muscle and depressor muscle, respectively. This was found within a limited frequency range.

<u>Details Simulation 2:</u> Corresponding simulations, now following the experiments of (Hess & Büschges, 1999), are shown in Fig. 8 in the same format as used in Fig. 7. In Fig. 8A, the sensory input (bottom row, dark blue line) was applied using shorter periods (3.6 s) compared to those of pure pilocarpine-driven periods (data as in Fig. 7A: 10.4 s, 4.6 s, 4.1 s, for alpha joint, beta joint and gamma joint, respectively). Therefore, as in the biological data, in all three joints pilocarpine-elicited oscillations are dominated by sensory influences. Fig. 8B shows the corresponding experiment but now with longer sensory period (7.0 s, bottom row, dark blue line). The alpha joint is still fully controlled by the sensory input, as the cycle remains shorter than the pilocarpine driven period (7.0 s vs.10.4 s). This is different for the beta joint (pilocarpine period: 4.6 s), and the same is true for the gamma joint (pilocarpine period: 4.1 s). In both these cases, influences from the pilocarpine-driven system can be seen. As in the biological experiments, reset effects can be observed in each joint. As a further effect, in both Fig. 8A and 8B, in the gamma joint the first half cycle (the pair of Flexor/Extensor) shows an inverted temporal sequence compared to later cycles. Thus, as shown in the earlier experiment (Fig. 7), sensory input may dominate slower pilocarpine-driven cycles, and resets the pilocarpine cycles if the latter is longer. To summarize: the simulation results show qualitatively the same results as observed in the biological experiments.

## 4 Discussion

In an earlier study (Schilling & Cruse, 2020), we followed the goal to find a minimal architecture called neuroWalknet, a simulator that is able to explain diverse behaviors of stick insects but might also in part be generalized to other insects. That study focused on behaviors concerning interleg control as are forward and backward walking, a broad range of velocities, various—including “non-canonical”—gaits, negotiation of curves, climbing on irregular ground, but also partial or complete deafferentation of legs. We used an embodied simulation (i.e., with a dynamically simulated or physical body), and introduced decentralized structures that enable control of the individual legs. In this current study, we now ask to what extent a number of influential experimental studies which specifically deal with *intraleg* control, could be explained by the control structure neuroWalknet, too. The studies in this article have in common that sensory stimulation in a leg directly causes a local reaction in the different joints. The studies cover stimulation of CS affecting the alpha (TC) joint (Akay et al., 2004, 2007; Schmitz, 1993), stimulation of femoral fCO affecting the beta (CT) joint (Hess & Büschges, 1999) or the gamma (FT) joint (Bässler, 1976, 1986, 1988; Bässler & Büschges, 1990; Bässler & Koch, 1989; Hellekes et al., 2012; Nothof & Bässler, 1990). As a further group of experiments, the influence of pilocarpine to the alpha joint or to the beta joint was studied. In some of these cases, reflex reversal is observed depending on different contexts as are forward – backward walking, passive and active states, or in specific situations of curve walking. To represent these states, three pairs of motivation units (Fig. 1) have been introduced to neuroWalknet as an extension and are used to control different contexts on different levels and thereby enable various combinations of neuronal structures to be activated.

To simulate all these experiments with neuroWalknet—as a general approach used in simulation studies—the structure is simplified compared to that of the biological system. In particular, the number of artificial neuronal units is much smaller than that of the biological counterpart, for there is only one instance of velocity unit (but see (Bender et al., 2010; Kai & Okada, 2013)), one unit for backward walking and for forward walking each (but see (Bidaye et al., 2014; DeAngelis et al., 2019; Feng et al., 2020)), and only two antagonistic motor units for each leg joint.

The introduced experiments required to add two specific sensory units to neuroWalknet, one for CS stimulation, the other for stimulation of the receptor apodeme of the fCO. As in the original neuroWalknet such sensors have been used only in a more qualitative way. In addition, we changed the value of the pilocarpine inputs, but kept the frequency values in the same range (see Methods). For each of the experiments addressed, we selected a qualitative, generalizable case, i.e., we focused on front legs.

As a result, the behavior found in all groups of biological experiments could be qualitatively replicated and explained by the simulation. As these have not been tested before in neuroWalknet, this further strengthens the idea that the structure of neuroWalknet represents essential aspects of the stick insect controller covering a broad range of hexapod walking, including the biological experiments simulated in the current article. The simulation results do not only refer to the biological experiments addressed here, but in addition predict data for all three joints (only one joint has been studied in the biological experiments). Thus, these additional data provide prediction for future studies and results.

### Specific phenomena as Active Reaction may be explained by a more general structure

Generally, these new results provide another example demonstrating that complex behavior does not necessarily require complex neuronal structures (see also Navinet (Cruse & Wehner, 2011; Hoinville & Wehner, 2018), positive feedback-based walking (Chittka & Niven, 2009; Schmitz et al., 2008)).

In particular, the intensively studied Active Reaction (Bässler, 1976, 1986, 1988; Bässler & Büschges, 1990) has previously led to the interpretation that the Active Reaction requires introducing an additional specific structure acting as positive feedback element that is used in specific situations. As a consequence, this interpretation led to simulation concepts as developed by Schmitz et al. (Schmitz et al., 2008) or Goldsmith et al. (Goldsmith et al., 2021). The former implemented a specific positive feedback solution without CPGs, whereas Goldsmith et al. (Goldsmith et al., 2021) built on the hypothesis that CPGs are required for walking. However, the current results offer another simpler interpretation in which results related to Active Reaction could be explained without requiring an additional neuronal structure. In the neuroWalknet approach, we attempted to realize Bässler’s (1988) postulate of a more general structure. This network embraces the whole leg with all three joints and allows for different behaviors as forward and backward walking, as well as—rather arhythmic—curve walking. In this alternative approach, neuroWalknet simply produces a velocity signal as assumed by different studies (e.g. (Cruse, 1985; Dean, 1984; Watson et al., 2002; Weiland & Koch, 1987)). This global feedforward velocity signal will then be controlled by states Walk, Forward or Backward, Stance or Swing and possibly, via walking direction “theta” by the ring net. This structure produces results that correspond to observations that can been called Active Reaction and includes an Active Reaction-like behavior also for the alpha joint ((Bässler, 1986), see Fig. 4A). Furthermore, in some situations no Active Reaction was observed in the animals in the biological experiment which agrees with our simulation results, too (e.g. Fig. 4B). As an example, during negotiation of curves, the ring net in neuroWalknet provides an Active Reaction-like movement for the inner leg, and for the outer leg. These are either a passive reflex-like behavior or a response that could be seen as a neutral reaction (between active and passive reflex). Our results (Fig. 3) correspond well to the biological experiments.

In other words, in this view Active Reaction does not represent a dedicated neuronal structure, but a phenomenon that eventually emerges in specific situations. Thus, Active Reaction as a separate neuronal element may only exist in the eye of the observer. What we earlier have seen as a separate structural element, may now be considered as an aspect of a more general holistic structure and an emergent property of this simpler structure.

The details as to how the information given by the sensory input might be transmitted within the neuronal system is still open. In a recent article, Goldsmith et al. (Goldsmith et al., 2021) proposed the idea, supported by their quantitative simulations of a leg model, that the desDUM neurons (Stolz et al., 2019) may provide a significant contribution and they further argue, based on studies of Sauer et al. (Sauer et al., 1996) and Driesang & Büschges (Driesang & Büschges, 1996), that a switch between states Walk and Stand on the local level may at least partly depend on changes of non-spiking interneurons.

### Open questions

An open point concerns the implementation of the neural structures that represents the effect of caudal bending (Akay et al., 2007; Schmitz, 1993), or the effect of ramp-like stimulation by elongation of the femoral fCO receptor apodeme (Bässler & Büschges, 1990; Hellekes et al., 2012), that lead to activation of units Swing or Stance. We did not try to install a specific neuronal structure but simply applied these signals to neuroWalknet (see the uppermost turquoise unit in Fig. 1, which in neuroWalknet corresponds to unit 170 and 171). For caudal bending, the corresponding sensors seem to be represented by several groups of trochanteral CS (Haberkorn et al., 2019). It is, however, even less known how exactly the ramp signal is transformed into an appropriate signal that activates state Stance. Correspondingly, the neuronal structures that connect stimulation of abdomen or antennae with unit FW or BW are unknown.

Another difference between legs has to be mentioned. In the experiments of Akay et al. (Akay et al., 2007) CS stimulation by bending the femur elicited retractor activation in forward walking and protractor activation in backward walking. This is different in the hind legs, but only when all other legs have been deafferented. This observation would support the idea (Nothof & Bässler, 1990) that the isolated hind leg has an inherent tendency to walk backwards, a concept not yet implemented in neuroWalknet.

### Central Pattern Generators

As shown in the results, figures 7 and 8, we simulated neuronal systems (Akay et al., 2007; Hess & Büschges, 1999) that were stimulated not only by sensory input (CS, fCO), but were in addition treated with pilocarpine, which leads to oscillations in the antagonistic structure of each joint. This is interesting because it is often assumed that such— postulated—CPGs form the basis to control quasi-rhythmic walking whereby these rhythmic elements are modulated by sensory input (e.g. Akay et al., p. 3285 and refs.), i.e., CPG activations are assumed to operate during walking even if no pilocarpine is applied.

Our simulations not using pilocarpine (Figures 2 - 5) show that such a CPG structure is structurally not necessary, as already demonstrated in our earlier study (Schilling & Cruse, 2020). Nonetheless, our current simulations (Figures 7, 8) demonstrate that the pilocarpine-driven experiments allow for oscillating motor output in neuroWalknet as well without an explicit CPG structure and that the structures in neuroWalknet that show oscillations driven by the pilocarpine input can be dominated by sensory input, or can effect reset, as shown in the biological experiments.

How might these results be explained? To this end, two hypothetical types of underlying structures should be compared that have been discussed in the literature. First, in a CPG based account, it is assumed that in each joint there exists an internal CPG structure that is used for walking and may also be activated when treated with pilocarpine. Such a structure may correspond to a combination of both structures shown in Fig. 6, with (a) inhibitory neurons equipped with phasic properties (as in Fig. 6, left, PMN), but in addition (b) a common input that, for example, may be connected with unit Walk, and even further (c) be connected with pilocarpine inputs. In comparison, in neuroWalknet as a second hypothetical network, there is no such internal CPG structure, but only the structure as shown in Fig. 6 (left, PMN), that (a) contains inhibitory neurons with phasic properties and (b), through external input from pilocarpine, may produce oscillations when activated. In this case, the internal structure as such does not allow for oscillations, but application of (simulated) pilocarpine (Fig. 6, left, purple arrows) may trigger the network to start oscillations. Thus, in this case the experiments shown in figures 7 and 8 could be explained without a CPG being necessary. Taking both hypotheses into account, on the one hand, the existence of CPGs used for walking could not be excluded for the first version, but, on the other hand, as shown by neuroWalknet, there is also no proof for the existence of such a CPG and these appear as not necessary. Note that, although the morphological differences between both such hypothetical networks are small, there is an overall functional difference. neuroWalknet appears to provide a functional advantage as no taming of quite a number of CPGs is required when stopping or when dealing with irregular substrate and negotiation of tight curves.

Importantly, our assumptions on neuroWalknet are not meant as a claim that there may no other CPGs being active in stick insects. The study of rocking behavior (Pflüger, 1977) strongly support the existence of CPGs being installed. Another suggestion comes from Bässler et al. (Bässler et al., 1991), who studied searching movements (for details see Supplement). They observed leg oscillations of up to 10 Hz, as well as similar data from Drosophila (Berendes et al., 2016), which also suggests the involvement of dedicated CPGs. Interestingly, Stoltz et al. (Stolz et al., 2019) studied influences of CS on stance activation of walking leg via desDUM neurons situated in the gnathal ganglion, but did not find influences of pilocarpine induced CPGs, which may support the idea that pilocarpine induced CPGs are not involved for control of slow walking. Similarly, Mentel et al. (Mentel et al., 2008) found DUM cells in the mesothoaric ganglion most of which were active during stance phase of the leg studied, but only one further unit (DUMna nl2) exhibited spontaneous rhythmic activity. However, this rhythm was not coupled to leg activity.

## 5 Conclusion

Taken together, neuroWalknet is considered a holistic approach that combines important behavioral elements within one complete system with the hope to better recognize the emergent properties of the system. We believe that such an approach leads to better understanding of the complete system, rather than considering only separate, local elements.

We further believe that neuroWalknet represents an RNN structure that combines the essential properties required for legged locomotion in insects (or, maybe even for arthropods). The basic structure consists of an RNN called motivation unit network (Stand-Walk, FW-BW, Stance-Swing) controlling sub-behaviors as are: Walk: curve, and swing or stance; Swing: swing movement and search; Stance: stance movement including height control, i.e. obstacle avoidance, negotiation of curves; Stand: The important case of standing behavior (addressed in the Supplement) dealing with negative position feedback including adaptive setpoint depending on substrate compliance is a future task.

Whereas some of these behaviors have been simulated, but not yet been integrated into neuroWalknet, the most relevant ones are represented by network structures that (i) allow for quantitative simulation of the behaviors, but may (ii) also serve as a scaffold to test new hypotheses concerning not yet considered behavior of alive animals, also with respect to other species. And (iii) allow the (simulated) robot to predict behaviors that have not yet been tested biologically (as, e.g., shown by data in dashed lines, Figs 2 – 8)).

Important, but still open questions refer to (i) a possible neuronal realization of the ring structure (see (Stone et al., 2017)), (ii) control of segregated hindlegs, (iii) proof of the existence of CPGs to be implemented during walking, and (iv) how to establish an appropriate solution for simulation of searching movements.

## 6 METHODS

We developed a simulation model in order to better understand the function of a complex system and to provide predictions that could be tested in later biological experiments (McClelland, 2009). Simulation models purposefully focus on specific, selected aspects and abstract away other details. A given model always introduces simplifications and every model is in one or the other sense “always wrong” (Box & Draper, 1987) as they focus on essential properties only (Kay, 2018). For instance, in our simulation the number of artificial neurons, for example motoneurons, is much smaller than that of the corresponding biological system. Likewise, the number of load sensors and their functional details are extremely simplified (Fig. 1, turquoise units) compared to those studied by (Haberkorn et al., 2019). Overall, models are a useful tool in computational neuroscience as they aim to summarize functional relations and aspects, provide explanations, and lead to falsifiable predictions (Kay, 2018).

The control model has been introduced in detail in (Schilling & Cruse, 2020) and we have explained extensions in section 2 and changes in this section. As a brief summary: The simulations consists of two main parts: On the one hand, the neuronal controller processing sensory inputs and producing control signals on a per leg basis (the neuroWalknet controller has been implemented in python (version 3), see https://github.com/hcruse/neuro_walknet and there the 2022 version). On the other hand, a dynamic simulation environment for the body of the hexapod robot Hector (Dürr et al., 2019; Schneider et al., 2014), which exists as a hardware version and as a dynamic simulation (implementations are publicly available: dynamical simulation environment is realized in C++ and based on the Open Dynamics Engine library, see https://github.com/malteschilling/hector). Here, we use the dynamic simulation.

The neural controller consists of artificial neurons. While the activation dynamics of neurons are often approximated using Hodgkin-Huxley differential equations (Hodgkin & Huxley, 1952), a reduced version has been used for simplification (Manoonpong et al., 2013).

The structure of neuroWalknet is decentralized: There is one identical control network for each of the six legs (see Fig. 1). Each of these leg controllers consists of three subnetworks (Fig. 1, bottom three rows, marked as alpha, gamma, beta), one for each joint. In addition, there are a number of global units that provide context information and allow to switch on or off walking and for selecting walking direction. Furthermore, there are units for interleg coordination realizing the coordination influence rules. On the motor level, the output is simplified as we consider for each joint two antagonistic muscles. For more details on neuroWalknet, see (Schilling & Cruse, 2020).

### Following the biological experiments, we had to adjust the sensory inputs in the simulation: In the

first and fourth simulated experiment (see Figs. 2 and 7) the animal is stimulated by (e.g., abdominal) body contact to adopt the walking state in general and by stimulation of specific campaniform sensilla (CS) of a specific leg to activate state swing or state stance. The detailed connectivity in the biological system is not yet known. In neuroWalknet (Fig. 1), this task has been simulated by body contact sensors that activate units Forward or Backward, which in turn activate unit Walk. In addition, stimulation of specific campaniform sensilla—via the turquoise units and units Swing or unit Stance—activate unit Walk, too.

Likewise, in the biological experiments—simulated in experiment two, three, and four, see figures 3, 4, 5, 8—specific ramp and hold stimulation of the fCO plus tactile body stimulation elicit specific motor output. Again, it is not known in detail how these connections are realized in the biological system, but a ramp stimulus with velocity faster than about 6 mm/s is assumed to lead to a resistance reflex whereas velocities slower than about 2 mm/s usually elicit assistance reflexes (Bässler, 1988). To simulate this in the network, in neuroWalknet the ramp-wise input is assumed to activate the following two channels (see Figures 3, 4, 5, 8). The first channel (dark blue, fCO) stimulates unit Stance as is done by the campaniform sensilla (see above). The second channel (light blue, fCOpos)—driven by fCO—, activates the height controller that also contributes to motor output (Fig. 1, for details see (Schilling & Cruse, 2020), Fig. 2, lower part, beta joint controller) as elongation of the fCO, illustrated by the ramp “FeTi flexion”, elicits levation of beta joint. Therefore, in figures 3, 4, 5, 8 the sensory input is represented by two fCO channels, fCO, (dark blue) that activates units stance or swing, and fCOpos (light blue), that activates position of the beta joint (i.e. the Flexor – Depressor system).

The second part of our simulation is constituted by the simulator for the animal behavior which is crucial as we are dealing with an embodied system (Brooks, 1991). Although often neglected, embodiment is a crucial entity as the physical body may provide an essential contribution to avoid unnecessarily complex neuronal structures. However, as a consequence, the physical properties of the model (e.g., size, inertia) usually differ from that of the biological system. In our case body length of the hexapod robot Hector (Dürr et al., 2019) is larger by a factor of about 10. Due to different inertia, the kind of movement may also be different: In our case walking velocity of robot Hector is about a factor of 10 slower (for more details see (Schilling & Cruse, 2020).

Another relevant aspect concerns the structural basis of so-called central pattern generators, being studied in a specific set of biological experiments where the neuronal network is treated with pilocarpine. In such experiments, usually sensory input and motor output channels and specific connectives are cut (Büschges et al., 1995). When treated with pilocarpine, the motor outputs of each of the three leg joints oscillate rhythmically, usually with a period in the range of 4 s to 10 s. These cycle frequencies are quite variable (they depend on pilocarpine concentration and probably on further parameters, as well as on the leg joint, see Table 2). As different studies show quite different values, in neuroWalknet we based our experiments on data given by (Büschges et al., 1995) (Table 2, 1^st^ line, ‘BSB’), as they provide the most solid data base and used similar periods in our simulation (Table 2, line 4, output pilo; for comparison: walking periods of *Carausius morosus* operate in the range of 0.5 s and 3 s (Graham, 1972)).

In the biological experiments used for comparison in experiment four, specific sensory inputs have been left intact. As a consequence, interesting interactions can be observed when sensory cycles take about equal the time of the induced oscillations or a bit faster than the cycles elicited by pilocarpine only (e.g. (Akay et al., 2007; Hess & Büschges, 1999)). To study this effect in simulation, we decided to apply artificial pilocarpine signals (Table 2, 3^rd^ line) with a frequency range corresponding to that of slow robot walking: Pilocarpine-driven cycles have been tuned to a period between 2 s and 10 s in the simulations (Table 2, 4^th^ line), i.e. these cycles are in the same range as we have used in earlier experiments (Schilling & Cruse, 2020), see Table 2, 7^th^ line. We selected input values in simulation that allowed some variability and cover a sensible range of periods. This was realized by selecting different sensory input patterns: these cyclic sensory stimuli (CS in Fig. 7, fCO in Fig. 8) consist of two parts, with a longer and a following shorter time window (the latter marked grey. Whereas in the earlier experiments not using pilocarpine (Figures 2–5) these time slots concern 4 s and 2 s respectively, i.e. a period of 6 s, in the experiments where pilocarpine is given (Figures 7, 8), different time slots for sensory stimulation have been tested and two interesting examples are shown (Fig. 7, period 3.6 s and 10.8 s; Fig. 8, period 3.6 s and 7.0 s Table 2 compares duration of the six motor output values, i.e. the period, for data elicited by pilocarpine only ((Büschges et al., 1995), Table 2, 1^st^ line, ‘BSB), and used for the current simulation ((Table 2, 4^th^ line ‘output pilo’). The corresponding periods are of about the same order in all three cases. However, the input values, i.e., the strength of the pilocarpine signal that elicits these periods (Table 2, 3^rd^ line), are smaller for the current version than those used in (Schilling & Cruse, 2020) (Table 2, 6^th^ line). This was chosen because higher input values could have dominated the effect of the sensory input.

## Acknowledgements

This research was supported by the research training group “DataNinja” (Trustworthy AI for Seamless Problem Solving: Next Generation Intelligence Joins Robust Data Analysis) funded by the German federal state of North Rhine-Westphalia and by the Cluster of Excellence Cognitive Interaction Technology CITEC (EXC 277) at Bielefeld University, which is funded by the German Research Foundation (DFG).

## References

Akay, T., Haehn, S., Schmitz, J., & Büschges, A. (2004). Signals from load sensors underlie interjoint coordination during stepping movements of the stick insect leg. Journal of Neurophysiology, 92.

Akay, T., Ludwar, B. C., Göritz, M. L., Schmitz, J., & Büschges, A. (2007). Segment Specificity of Load Signal Processing Depends on Walking Direction in the Stick Insect Leg Muscle Control System. Journal of Neuroscience, 27(12), 3285–3294. https://doi.org/10.1523/JNEUROSCI.5202-06.2007

Bässler, U. (1976). Reversal of a reflex to a single motoneuron in the stick insect Carausius morosus. Biological Cybernetics, 24, 47–49.

Bässler, U. (1983). *Neural basis of elementary behavior in stick insects*. Springer, Berlin, Heidelberg, New York.

Bässler, U. (1986). Afferent control of walking movements in the stick insect Cuniculina impigra. Journal of Comparative Physiology [A], 158, 351–362.

Bässler, U. (1988). Functional Principles of Pattern Generation for Walking Movements of Stick Insect Forelegs: The Role of the Femoral Chordotonal Organ Afferences. Journal of Experimental Biology, 136, 125–147.

Bässler, U. (1993). The walking-(and searching-) pattern generator of stick insects, a modular system composed of reflex chains and endogenous oscillators. Biological Cybernetics, 69(4), 305–317. https://doi.org/10.1007/BF00203127

Bässler, U., & Büschges, A. (1990). Interneurones Participating in the "Active Reaction" in Stick Insects. Biological Cybernetics, 62, 529–538.

Bässler, U., & Koch, U. T. (1989). Modelling of the active reaction of stick insects by a network of neuromimes. Biological Cybernetics, 62(2), 141–150. https://doi.org/10.1007/BF00203002

Bässler, U., Rohrbacher, J., Karg, G., & Breutel, G. (1991). Interruption of searching movements of partly restrained front legs of stick insects, a model situation for the start of a stance phase? Biological Cybernetics, 65(6), 507–514. https://doi.org/10.1007/BF00204664

Bender, J. A., Pollack, A. J., & Ritzmann, R. E. (2010). Neural activity in the central complex of the insect brain is linked to locomotor changes. Curr. Biol, 20, 921–926.

Berendes, V., Zill, S. N., Bockemühl, B. A., & T. (2016). Speed-dependent interplay between local pattern-generating activity and sensory signals during walking in Drosophila. Journal of Experimental Biology, 219, 3781–3793. https://doi.org/10.1242/jeb.146720

Berg, E., Hooper, S. L., Schmidt, J., & Büschges, A. (2015). A Leg-Local Neural Mechanism Mediates the Decision to Search in Stick Insects. Current Biology, 25. https://doi.org/10.1016/j.cub.2015.06.017

Bidaye, S. S., Machacek, C., Wu, Y., & Dickson. (2014). Neuronal Control of Drosophila Walking Direction. Science, 344, 97–101.

Bläsing, B. (2006). Crossing Large Gaps: A Simulation Study of Stick Insect Behavior. Adaptive Behavior, 14(3), 265–285.

Bläsing, B., & Cruse, H. (2004). Stick insect locomotion in a complex environment: Climbing over large gaps. Journal of Experimental Biology, 207, 1273–1286. https://doi.org/10.1242/jeb.00888.

Box, G. E. P., & Draper, N. R. (1987). Empirical model-building and response surfaces (pp. xiv, 669). John Wiley & Sons.

Brooks, R. A. (1991). Intelligence without representation. Artificial Intelligence, 47, 139–159.

Bucher, D., Akay, T., DiCaprio, R. A., & Büschges, A. (2003). Interjoint coordination in the stick insect leg-control system: The role of positional signaling. Journal of Neurophysiology, 89(3), 1245–1255.

Büschges, A., Schmitz, J., & Bässler, U. (1995). Rhythmic patterns in the thoracic nerve cord of the stick insect induced by pilocarpine. J. Exp. Biol, 198,435*-*456.

Chittka, L., & Niven, J. (2009). Are Bigger Brains Better? Current Biology, 19(21), R995–R1008. https://doi.org/10.1016/j.cub.2009.08.023

Cruse, H. (1985). Which parameters control the leg movement of a walking insect? I. Velocity control during the stance phase. The Journal of Experimental Biology, 116, 343--355.

Cruse, H., Kindermann, T., Schumm, M., Dean, J., & Schmitz, J. (1998). Walknet—A biologically inspired network to control six-legged walking. Neural Netw, 11(7–8), 1435–1447. http://dx.doi.org/10.1016/S0893-6080(98)00067-7

Cruse, H., Kühn, S., Park, S., & Schmitz, J. (2004). Adaptive control for insect leg position: Controller properties depend on substrate compliance. Journal of Comparative Physiology A, 190(983--991).

Cruse, H., & Wehner, R. (2011). No Need for a Cognitive Map: Decentralized Memory for Insect Navigation. PLoS Comput. Biol, 7(3), 1002009.

Dean, J. (1984). Control of leg protraction in the stick insect: A targeted movement showing compensation for externally applied forces. Journal of Comparative Physiology A, 155(6), 771–781. https://doi.org/10.1007/BF00611594

DeAngelis, B. D., Zavatone-Veth, J. A., & Clark, D. A. (2019). The manifold structure of limb coordination in walking Drosophila. ELife, 8, e46409. https://doi.org/10.7554/eLife.46409

Driesang, R. B., & Büschges, A. (1996). Physiological changes in central neuronal pathways contributing to the generation of a reflex reversal. Journal of Comparative Physiology A, 179(1), 45–57.

Dürr, V. (2001). Stereotypic leg searching movements in the stick insect: Kinematic analysis, behavioural context and simulation. Journal of Experimental Biology, 204, 1589–1604.

Dürr, V., Arena, P., Cruse, H., Dallmann, C. J., Drimus, A., Hoinville, T., Krause, T., Mátéfi-Tempfli, S., Paskarbeit, J., Patanè, L., Schäffersmann, M., Schilling, M., Schmitz, J., Strauss, R., Theunissen, L., Vitanza, A., & Schneider, A. (2019). Integrative Biomimetics of Autonomous Hexapedal Locomotion. Frontiers in Neurorobotics, 13. https://doi.org/10.3389/fnbot.2019.00088

Dürr, V., & Ebeling, W. (2005). The behavioural transition from straight to curve walking: Kinetics of leg movement parameters and the initiation of turning. Journal of Experimental Biology, 208(12), 2237–2252.

Dürr, V., Schmitz, J., & Cruse, H. (2004). Behaviour-based modelling of hexapod locomotion: Linking biology and technical application. Arthropod Structure \& Development, 33(3), 237– 250.

Ebeling, W., & Dürr, V. (2006). Perturbation of leg protraction causes context-dependent modulation of inter-leg coordination, but not of avoidance reflexes. Journal of Experimental Biology, 209(11), 2199–2214.

Ekeberg, O., Blümel, M., & Büschges, A. (2004). Dynamic simulation of insect walking. Arthropod Structure \& Development, 33(3), 287–300.

Feng, K., Sen, R., Minegishi, R., Dübbert, M., Bockemühl, T., Büschges, A., & Dickson, B. J. (2020). Distributed control of motor circuits for backward walking in Drosophila. Nature Communications, 11(1), 1–17. https://doi.org/10.1038/s41467-020-19936-x

Goldsmith, C. A., Quinn, R. D., & Szczecinski, N. S. (2021). Investigating the role of low level reinforcement reflex loops in insect locomotion. Bioinspiration & Biomimetics, 16(6), 065008. https://doi.org/10.1088/1748-3190/ac28ea

Graham, D. (1972). A behavioural analysis of the temporal organisation of walking movements in the 1st instar and adult stick insect. J.Comp.Physiol, 81,*23*, 52.

Haberkorn, A., Gruhn, M., Zill, S. N., & Büschges, A. (2019). Identification of the origin of force-feedback signals influencing motor neurons of the thoraco-coxal joint in an insect. Journal of Comparative Physiology A, 205(2), 253–270.

Hellekes, K., Blincow, E., Hoffmannn, J., & Büschges, A. (2012). Control of reflex reversal in stick insect walking: Effects of intersegmental signals, changes in direction, and optomotor-induced turning. J. Neurophysiol, 107, 239–249.

Hess, D., & Büschges, A. (1999). Role of Proprioceptive Signals From an Insect Femur-Tibia Joint in Patterning Motoneuronal Activity of an Adjacent Leg Joint. Journal of Neurophysiology, 81(4), 1856–1865. https://doi.org/10.1152/jn.1999.81.4.1856

Hodgkin, A. L., & Huxley, A. F. (1952). A quantitative description of membrane current and its application to conduction and excitation in nerve. J. Physiol, 117, 500–544.

Hoinville, T., & Wehner, R. (2018). Optimal multiguidance integration in insect navigation. Proceedings of the National Academy of Sciences, 115(11), 2824–2829. https://doi.org/10.1073/pnas.1721668115

Kai, K., & Okada, J. (2013). Characterization of Locomotor-Related Spike Activity in Protocerebrum of Freely Walking Cricket. Zoological Science, 30, 591–601.

Kay, K. N. (2018). Principles for models of neural information processing. NeuroImage, 180, 101– 109. https://doi.org/10.1016/j.neuroimage.2017.08.016

Lévy, J., & Cruse, H. (2008a). Controlling a system with redundant degrees of freedom. I. Torque distribution in still standing stick insects. *Journal of Comparative Physiology A: Neuroethology, Sensory*, Neural, and Behavioral Physiology, 194(8), 719--733. https://doi.org/10.1007/s00359-008-0343-1

Lévy, J., & Cruse, H. (2008b). Controlling a system with redundant degrees of freedom. II. Solution of the force distribution problem without a body model. *Journal of Comparative Physiology A: Neuroethology, Sensory*, Neural, and Behavioral Physiology, 194(8), 735--750. https://doi.org/10.1007/s00359-008-0343-1

Manoonpong, P., Parlitz, U., & Wörgötter, F. (2013). Neural control and adaptive neural forward models for insect-like, energy-efficient, and adaptable locomotion of walking machines. Frontiers in Neural Circuits. https://doi.org/10.3389/fncir.2013.00012

McClelland, J. L. (2009). The Place of Modeling in Cognitive Science. Topics in Cognitive Science, 1(1), 11–38. https://doi.org/10.1111/j.1756-8765.2008.01003.x

Mentel, T., Weiler, V., Büschges, A., & Pflüger, H.-J. (2008). Activity of neuromodulatory neurones during stepping of a single insect leg. Journal of Insect Physiology, 54(1), 51–61. https://doi.org/10.1016/j.jinsphys.2007.08.010

Nothof, U., & Bässler, U. (1990). The network producing the “active reaction” of stick insects is a functional element of different pattern generating systems. Biological Cybernetics, 62(5), 453–462.

Pflüger, H.-J. (1977). The control of the rocking movements of the phasmidCarausius morosus Br. Journal of Comparative Physiology, 120(2), 181–202. https://doi.org/10.1007/BF00619314

Pflüger, H.-J., Büschges, A., & Bässler, U. (2021). Historical Review on Thanatosis with Special Reference to the Work of Fritz Steiniger. In M. Sakai (Ed.), Death-Feigning in Insects: Mechanism and Function of Tonic Immobility (pp. 15–21). Springer. https://doi.org/10.1007/978-981-33-6598-8_2

Rosano, H., & Webb, B. (2007). A dynamic model of thoracic differentiation for the control of turning in the stick insect. Biological Cybernetics, 97(3), 229--246.

Sauer, A. E., Driesang, R. B., Büschges, A., Bässler, U., & Borst, A. (1996). Distributed processing on the basis of parallel and antagonistic pathways simulation of the femur-tibia control system in the stick insect. Journal of Computational Neuroscience, 3(3), 179–198.

Schilling, M., & Cruse, H. (2017). ReaCog, a Minimal Cognitive Controller Based on Recruitment of Reactive Systems. Frontiers in Neurorobotics, 11. https://pub.uni-bielefeld.de/record/2908308

Schilling, M., & Cruse, H. (2020). Decentralized control of insect walking: A simple neural network explains a wide range of behavioral and neurophysiological results. PLOS Computational Biology, 16(4), e1007804. https://doi.org/10.1371/journal.pcbi.1007804

Schilling, M., Hoinville, T., Schmitz, J., & Cruse, H. (2013). Walknet, a bio-inspired controller for hexapod walking. Biol Cybern, 107(4), 397–419. https://doi.org/10.1007/s00422-013-0563-5

Schilling, M., Konen, K., Ohl, F. W., & Korthals, T. (2020). Decentralized Deep Reinforcement Learning for a Distributed and Adaptive Locomotion Controller of a Hexapod Robot. 8.

Schilling, M., & Melnik, A. (2018). An Approach to Hierarchical Deep Reinforcement Learning for a Decentralized Walking Control Architecture. Biologically Inspired Cognitive Architectures 2018. Proceedings of the Ninth Annual Meeting of the BICA Society, 848. https://pub.uni-bielefeld.de/record/2934190

Schilling, M., Melnik, A., Ohl, F. W., Ritter, H. J., & Hammer, B. (2021). Decentralized control and local information for robust and adaptive decentralized Deep Reinforcement Learning. Neural Networks, 144, 699–725. https://doi.org/10.1016/j.neunet.2021.09.017

Schilling, M., Paskarbeit, J., Hoinville, T., Hüffmeier, A., Schneider, A., Schmitz, J., & Cruse, H. (2013). A hexapod walker using a heterarchical architecture for action selection. Front Comput Neurosci, 7, 126. https://doi.org/10.3389/fncom.2013.00126

Schilling, M., Paskarbeit, J., Ritter, H., Schneider, A., & Cruse, H. (2021). From Adaptive Locomotion to Predictive Action Selection – Cognitive Control for a Six-Legged Walker. IEEE Transactions on Robotics, 1–17. https://doi.org/10.1109/TRO.2021.3106832

Schmitz, J. (1993). Load compensatory reactions in the proximal leg joints of stick insects during standing and walking. Journal of Experimental Biology, 183, 15–33.

Schmitz, J., Schneider, A., Schilling, M., & Cruse, H. (2008). No need for a body model: Positive velocity feedback for the control of an 18-DOF robot walker. *Applied Bionics and Biomechanics, Special Issue on Biologically Inspired Robots*, *5*(3), 135–147.

Schmitz, J., & Stein, W. (2000). Convergence of load and movement information onto leg motoneurons in insects. Journal of Neurobiology, 42(4), 424–436.

Schneider, A., Cruse, H., & Schmitz, J. (2006). Decentralized Control of Elastic Limbs in Closed Kinematic Chains. The International Journal of Robotics Research, 25(9), 913--930.

Schneider, A., Paskarbeit, J., Schilling, M., & Schmitz, J. (2014). HECTOR, a bio-inspired and compliant hexapod robot. Proceedings of the 3rd Conference on Biomimetics and Biohybrid Systems.

Stolz, T., Diesner, M., Neupert, S., Hess, M. E., Delgado-Betancourt, E., Pflüger, H.-J., & Schmidt, J. (2019). Descending octopaminergic neurons modulate sensory-evoked activity of thoracic motor neurons in stick insects. Journal of Neurophysiology, 122(6), 2388–2413.

Stone, T., Webb, B., Adden, A., Weddig, N. B., Honkanen, A., Templin, R., Wcislo, W., Scimeca, L., Warrant, E., & Heinze, S. (2017). An Anatomically Constrained Model for Path Integration in the Bee Brain. Current Biology, 27, 3069–3085. https://doi.org/10.1016/j.cub.2017.08.052

Watson, J., Ritzmann, R., & Pollack, A. (2002). Control of climbing behavior in the cockroach, Blaberus discoidalis. II. Motor activities associated with joint movement. *Journal of Comparative Physiology A: Sensory*, Neural, and Behavioral Physiology, 188(1), 55–69. https://doi.org/10.1007/s00359-002-0278-x

Weiland, G., & Koch, U. T. (1987). Sensory feedback during active movements of stick insects. Journal of Experimental Biology, 133(1), 137–156.

